# Plxnd1-mediated mechanosensing of blood flow controls the caliber of the Dorsal Aorta via the transcription factor Klf2

**DOI:** 10.1101/2024.01.24.576555

**Authors:** Jia He, Adriana Blazeski, Uthayanan Nilanthi, Javier Menéndez, Samuel C. Pirani, Daniel S. Levic, Michel Bagnat, Manvendra K. Singh, José G Raya, Guillermo García-Cardeña, Jesús Torres-Vázquez

## Abstract

The cardiovascular system generates and responds to mechanical forces. The heartbeat pumps blood through a network of vascular tubes, which adjust their caliber in response to the hemodynamic environment. However, how endothelial cells in the developing vascular system integrate inputs from circulatory forces into signaling pathways to define vessel caliber is poorly understood. Using vertebrate embryos and *in vitro*-assembled microvascular networks of human endothelial cells as models, flow and genetic manipulations, and custom software, we reveal that Plexin-D1, an endothelial Semaphorin receptor critical for angiogenic guidance, employs its mechanosensing activity to serve as a crucial positive regulator of the Dorsal Aorta’s (DA) caliber. We also uncover that the flow-responsive transcription factor KLF2 acts as a paramount mechanosensitive effector of Plexin-D1 that enlarges endothelial cells to widen the vessel. These findings illuminate the molecular and cellular mechanisms orchestrating the interplay between cardiovascular development and hemodynamic forces.

**Highlights:**

- Plexin-D1 mechanosensing of blood flow tunes the caliber of the Dorsal Aorta (DA)
- The DA widens without raising endothelial cell numbers, which can change separate from the caliber
- The Kruppel-like transcription factor 2 (KLF2) is a key Plexin-D1 mechano-effector during development
- KLF2 increases endothelial cell size to expand the DA caliber

## INTRODUCTION

Genetic programs and physical forces interact to control signaling pathways, gene expression, and cellular behavior. Their interplay directs embryonic morphogenesis and organ formation, function, and homeostasis. A prominent arena where this coordination plays out is the cardiovascular system, where blood flow forces impact the development of the heart and blood vessels and vice versa. The circulatory function of the cardiovascular system relies on three main factors: the heart’s pumping activity, the hierarchical branching of blood vessels, and their ability to adapt their caliber or luminal size to fluid forces and tissue demands[1-5].

According to Poiseuille’s law, vascular resistance depends on vessel caliber, length, and blood viscosity. Vascular resistance is the opposition the heart’s contractions must overcome to propel blood through vessels and establish circulatory flow. Its value is inversely proportional to the fourth power of the blood vessel’s radius. Therefore, the impact of slight alterations in vessel caliber on vascular resistance can be noteworthy. To illustrate, reducing a vessel’s luminal radius by ten percent increases its vascular resistance by fifty percent[6].

Vessel caliber influences hemodynamics (blood pressure and tissue perfusion), vascular remodeling, vessel integrity, and the structure and function of the heart. Thus, vessel caliber exerts a global impact on the cardiovascular system’s development, performance, and health. Vascular caliber abnormalities lead to cardiovascular diseases that can culminate in disability or death[6-20].

Blood circulation generates mechanical forces that act on the endothelial vascular wall, including circumferential, axial, and shear stress. Circumferential and axial stress depend on intraluminal pressure and act around the vessel’s perimeter or length. In contrast, shear stress derives from the friction between blood flow and the endothelial cells (ECs) lining the vessel and occurs parallel to its wall. The magnitude of shear stress primarily depends on the vascular caliber, with secondary influences from the flow rate and the blood’s viscosity[6, 21, 22].

ECs express specialized proteins that sense the mechanical forces induced by the circulation. The transformation of these mechanical inputs into intracellular biochemical responses enables blood vessels to adapt to varying hemodynamic conditions using different strategies. These include changes in vessel caliber, the acquisition of an atheroprotective endothelial gene expression profile that suppresses blood clotting and inflammation, and modifications in endothelial barrier function[23-25].

Mammalian Plexin-D1 (Plxnd1 in the zebrafish) is a Semaphorin receptor specific to vertebrates located on the cell surface of various tissues. In the endothelium, this transmembrane protein plays a prominent and evolutionarily conserved role in guiding the stereotypical branching pattern of blood vessels[26-28]. Recent structural analyses and studies with cultured ECs have revealed that Plexin-D1 also functions as a mechanosensor of fluid shear stress[29].

However, the developmental role of the receptor’s novel mechanosensing function is unknown. To unveil it, we analyzed the impact of the guidance and mechanosensing activities of Plxnd1, along with blood flow, on the formation of the zebrafish embryonic cardiovascular system. Our studies reveal that Plxnd1 is a crucial component of the endothelial genetic circuit responsible for enabling the Dorsal Aorta (DA), one of the body’s largest arteries, to enlarge its caliber in response to increasing circulatory flow forces. Notably, the role of Plxnd1 in regulating DA caliber is distinct from its GAP (GTPase Activating Protein)-mediated guidance function.

Accordingly, *plxnd1* null fish show a decreased DA caliber. *KLF2* (*Krüppel-like Factor 2*) orthologs display fluid shear stress-inducible expression and encode atheroprotective transcription factors with emerging roles in vascular sizing[30-38]. We found that the DA’s circulation-induced endothelial upregulation of *klf2* levels is *plxnd1* dependent; see[29]. Furthermore, loss and gain-of-function experiments identify Klf2 as a pivotal Plxnd1 mechano-effector responsible for driving vascular caliber expansion by increasing the size of ECs. Moreover, elevating Klf2 expression in ECs can widen the narrow caliber of the DA in fish that lack blood flow, providing evidence that Klf2 activity *per se* can restore the effects of lack of blood flow on the caliber of DA.

Notably, additional experiments argue that the mechanisms by which the receptor regulates the DA’s caliber are similar in fish and mammals. First, murine embryos with endothelial *Plxnd1* deletion also display a narrow DA. Second, studies with perfusable microvascular networks engineered with human endothelial cells show that, like in the zebrafish, circulatory flow increases *KLF2* expression, expands vascular caliber, and enlarges endothelial cells.

Our findings expand our comprehension of the pivotal roles played by Plexin-D1 and KLF2 in cardiovascular development. They also predict the involvement of variants in these genes in diseases promoting vascular caliber abnormalities, such as atherosclerosis, cardiac ischemia, diabetes, hypertension, peripheral artery disease, Raynaud’s disease, retinopathy, and systemic sclerosis. Conversely, controlling the activity of these proteins activities could prevent or mitigate these conditions[39-43].

## RESULTS

### Plxnd1 acts in the endothelium with circulation-dependency as a positive regulator of the DA caliber

In the WT zebrafish embryo, the heart has not yet started beating at 24 hours post-fertilization (hpf), resulting in the absence of circulatory flow at this stage. Soon after, the heartbeat begins, and cardiac output and circulatory flow progressively increase[44-51]. In WTs, the trunk’s axial blood vessels respond to increased circulatory flow bi-phasically. They expand up to 48 hpf, narrowing afterward by 72 hpf[44, 45, 52].To evaluate the influence of circulatory flow and Plxnd1 on vascular caliber, we manipulated blood flow and Plxnd1 activities by utilizing recessive alleles. To visualize the vessels, we used the cytosolic endothelial reporter *Tg(fli1:EGFP)^y1^*[53]. Since the axial vessels are not perfect cylinders, we manually measured their cross-sectional luminal area to establish their caliber. We performed these quantifications before the circulation started (24 hpf) and after it had begun (32, 48, and 72 hpf).

To eliminate blood flow, we used *silent heart* (*sih*) mutants, which lack the heartbeat due to loss of *tnnt2a* (*troponin T type 2a (cardiac)*) expression (*sih^tc300b^*)[54]. The *tnnt2a* gene encodes a critical contractile myofibrillar component selectively expressed in the myocardium and the smooth muscle cells around the heart’s outflow tract. Notably, *sih* mutants show properly patterned angiogenic vessels (**Figures S1A, B**) up to 72 hpf and die around one week of age[54-57].

We manipulated *plxnd1* activity using homozygotes of four *plxnd1* alleles or WT embryos with morpholino (MO)-mediated *plxnd1* knockdown (*plxnd1* morphants). We utilized two chemically-induced *plxnd1* alleles to deactivate the receptor’s functions completely. The *plxnd1^Df(Chr08)fs31l^* allele is a homozygous-lethal chromosomal deficiency removing *plxnd1* and nearby genes. In contrast, *plxnd1^fov01b^* harbors a premature stop codon early within the receptor’s extracellular Sema domain. Its homozygotes show reduced *plxnd1* mRNA levels, suggesting mutant message degradation and translational suppression[58-60]. To selectively impair the receptor’s GAP (GTPase Activating Protein) function essential for its guidance activity, we used genome editing[61] to make *plxnd1^GAP1^* and *plxnd1^GAP2^* alleles (**Table S1**) by altering the position of the catalytic and stabilizing Arginine residues at the GAP1 and GAP2 cytosolic tail motifs[26, 62-64], respectively. These four mutants and the morphants display misguided angiogenic vessels (**Figures S1A, C-G**), robust circulation (**Figure S2**; see[58, 59]), and survive to adulthood (except for the *plxnd1^Df(Chr08)fs31l^*fish and *plxnd1* morphants). Considering the *plxnd1^Df(Chr08)fs31l^* allele’s lethal nature[58] and the phenotypic similarities between both *plxnd1^GAP^* mutants, we primarily relied on the *plxnd1^fov01b^* and *plxnd1^GAP1^* alleles for our analyses. Our initial studies revealed that circulatory flow and *plxnd1* might interact differently to dictate the caliber of each axial vessel. Here, we focus on dissecting their role in regulating the DA’s caliber.

We observed that the DA calibers of the WT-like siblings (WT-like sibs) of the *plxnd1^fov01b^*, *plxnd1^GAP1^*, and *sih* mutants are identical at 24 hpf when circulation has not yet begun (**Figures 1A-C**), indicating that DA lumenogenesis is independent of *plxnd1* and *tnnt2a* activities and circulatory flow. Afterward, concurrent with the increasing circulatory flow, we observed that in the mutant’s WT-like sibs, the DA caliber gradually expanded to 48 hpf and diminished by 72 hpf (**Figures 1A-C**), consistent with prior findings[52, 65].

**Figure 1.**
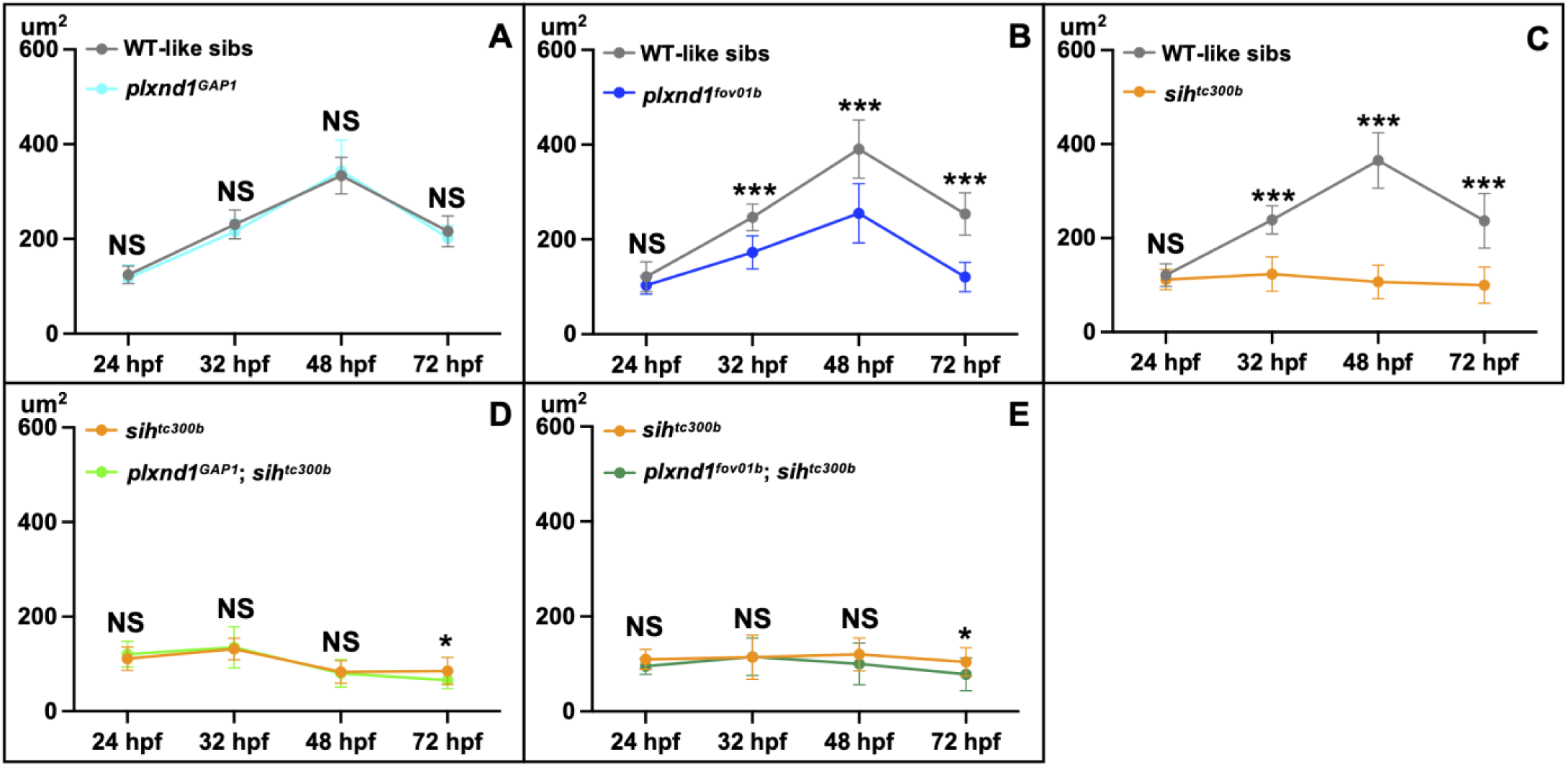
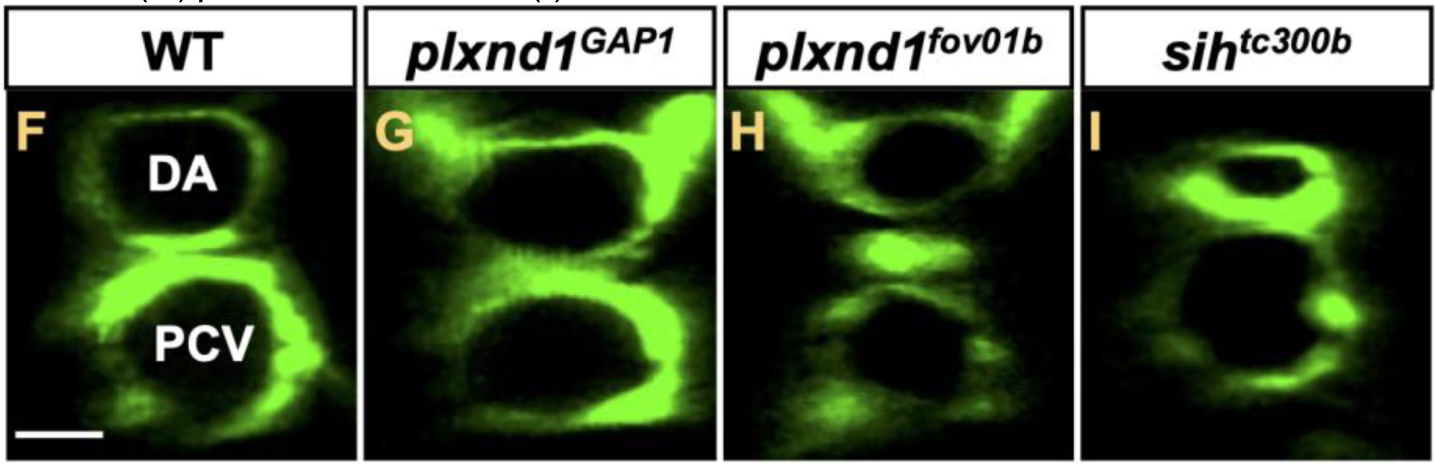
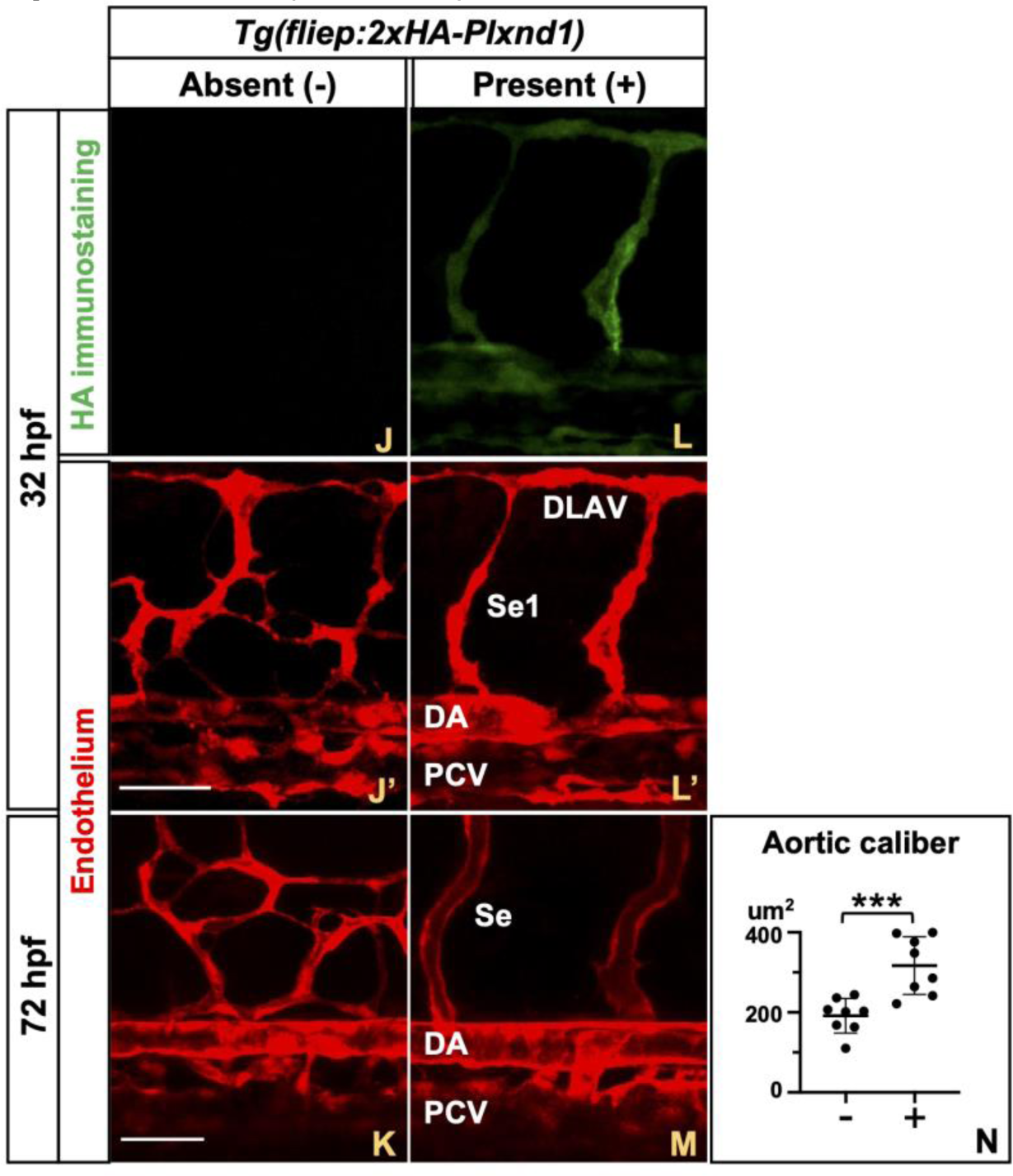
Endothelial Plxnd1 controls the caliber of the aorta in a circulation-dependent fashion. (**A-E**) Comparisons of aortic caliber (luminal area of aortic cross-sections) at 24, 32, 48, and 72 hpf in fixed siblings (sibs) of various genotypes. <u>Statistical measures</u> <u>are as follows</u>. The dots represent means. The error bars denote the standard deviation (SD). Significance levels as \**p*≤0.05, ∗∗*p*≤0.01, ∗∗∗*p*≤0.001; unpaired two-tailed Student’s *t*-test. (**A**) Comparison of WT-like sibs (grey) vs. *plxnd1^GAP1^* mutants (cyan). WT-like sibs group: WTs and both single (*plxnd1^GAP1^/+* and *sih^tc300b^/+*) and double (*plxnd1^GAP1^/+*; *sih^tc300b^/+*) heterozygotes. *plxnd1^GAP1^* mutant group: *plxnd1^GAP1^* and *plxnd1^GAP1^*; *sih^tc300b^/+*. WT-like sibs per stage: 24 hpf (n = 10), 32 hpf (n = 11), 48 hpf (n = 12), 72 hpf (n = 10). *plxnd1^GAP1^* mutants per stage: 24 hpf (n = 19), 32 hpf (n = 12), 48 hpf (n = 19), 72 hpf (n = 24). Significance levels per stage: 24 hpf (*p*=0.4936), 32 hpf (*p*=0.2120), 48 hpf (*p*=0.6335), 72 hpf (*p*=0.1840). (**B**) Comparison of WT-like sibs (grey) vs. *plxnd1^fov01b^* mutants (navy blue). WT- like sibs group: WTs and both single (*plxnd1^fov01b^/+* and *sih^tc300b^/+*) and double (*plxnd1^fov01b^/+*; *sih^tc300b^/+*) heterozygotes. *plxnd1^fov01b^* mutant group: *plxnd1^fov01b^* and *plxnd1^fov01b^*; *sih^tc300b^/+*. WT-like sibs per stage: 24 hpf (n = 12), 32 hpf (n = 11), 48 hpf (n = 11), 72 hpf (n = 13). *plxnd1^fov01b^* mutants per stage: 24 hpf (n = 15), 32 hpf (n = 10), 48 hpf (n = 15), 72 hpf (n = 16). Significance levels per stage: 24 hpf (*p*=0.2145), 32 hpf (*p*<0.0001), 48 hpf (*p*<0.0001), 72 hpf (*p*<0.0001). (**C**) Comparison of WT-like sibs (grey) vs. *sih^tc300b^* mutants (orange). WT-like sibs group: WTs and both single (*sih^tc300b^/+*, *plxnd1^fov01b^/+*, and *plxnd1^GAP1^/+*) and double (*plxnd1^fov01b^/+*; *sih^tc300b^/+* and *sih^tc300b^/+*; *plxnd1^GAP1^/+*) heterozygotes. *sih^tc300b^* mutant group: *sih^tc300b^* homozygotes alone and with *plxnd1* heterozygosity (*sih^tc300b^*; *plxnd1^GAP1^/+* and *sih^tc300b^*; *plxnd ^fov01b^/+*). WT-like sibs per stage: 24 hpf (n = 12), 32 hpf (n = 22), 48 hpf (n = 27), 72 hpf (n = 26). *sih^tc300b^* mutants per stage: 24 hpf (n = 13), 32 hpf (n = 21), 48 hpf (n = 27), 72 hpf (n = 22). Significance levels per stage: 24 hpf (*p*=0.3077), 32 hpf (*p*<0.0001), 48 hpf (*p*<0.0001), 72 hpf (*p*<0.0001). (**D**) Comparison of *sih^tc300b^* sibs (orange) vs. *plxnd1^GAP1^; sih^tc300b^* double mutants (light green). *sih^tc300b^* sibs group: *sih^tc300b^* and *sih^tc300b^*; *plxnd1^GAP1^/+*. The genotype of the double mutant group is self-explanatory. *sih^tc300b^* sibs per stage: 24 hpf (n = 7), 32 hpf (n = 11), 48 hpf (n = 10), 72 hpf (n = 12). *plxnd1^GAP1^; sih^tc300b^* double mutants per stage: 24 hpf (n = 15), 32 hpf (n = 7), 48 hpf (n = 15), 72 hpf (n = 15). Significance levels per stage: 24 hpf (*p*=0.4236), 32 hpf (*p*=0.8098), 48 hpf (*p*=0.7940), 72 hpf (*p*=0.0419). (**E**) Comparison of *sih^tc300b^* sibs vs. *plxnd1^fov01b^; sih^tc300b^* double mutants (dark green). *sih^tc300b^* sibs group: *sih^tc300b^* and *sih^tc300b^*; *plxnd1 ^fov01b^/+*. The genotype of the double mutant group is self-explanatory. *sih^tc300b^*sibs per stage: 24 hpf (n = 5), 32 hpf (n = 10), 48 hpf (n = 17), 72 hpf (n = 15). *plxnd1^fov01b^; sih^tc300b^* double mutants per stage: 24 hpf (n = 6), 32 hpf (n = 12), 48 hpf (n = 14), 72 hpf (n = 25). Significance levels per stage are as follows: 24 hpf (*p*=0.2434), 32 hpf (*p*=0.9735), 48 hpf (*p*=0.1637), 72 hpf (*p*=0.0221). (**F-I**) Representative DA and PCV cross-sections from live 48 hpf embryos reconstructed from confocal microscopy images. Endothelium (green), *Tg(fli1:EGFP)^y1^*. The dorsal side is up, and the left side is left. (**F**) WT. The scale bar (horizontal white line) is 15 μm. (**G**) *plxnd1^GAP1^* mutant. (**H**) *plxnd1^fov01b^* mutant. (**I**) *sih^tc300b^* mutant. (**J-N**) Forced endothelial expression of HA-tagged Plxnd1 rescues the angiogenic misguidance and the reduced DA caliber of *plxnd1^fov01b^*mutants. (**J-M**) Lateral views of the trunk of fixed sibling *plxnd1^fov01b^* mutants without (J-K) or with (L-M) the *Tg(fliep:2xHA-plxnd1)* transgene at 32 hpf (J-J’, L-L’) and live embryos 72 hpf (K, M). The dorsal side is up, and the anterior side is left. HA immunostaining, green. Endothelium (red), *Tg(kdrl:mCherry)^y206^* reporter[185]. The scale bar (horizontal white line) is 40 μm (J’, K). (**N**) Quantification of DA caliber in sibling *plxnd1^fov01b^* mutants at 72 hpf as a function of the absence (-) or presence (+) of the *Tg(fliep:2xHA-plxnd1)* transgene. Statistical measures. The dots represent individual data points. The horizontal lines represent the means. The error bars denote the SD. Significance level, \*\*\**p*≤0.001, unpaired two-tailed Student’s *t*-test.

The DA caliber of *plxnd1^GAP1^*mutants adjusts appropriately with changes in blood flow (**Figures 1A, F, G**). Likewise, *plxnd1^GAP2^* fish display the correct DA caliber in the presence of the circulation (not shown). However, *plxnd1^fov01b^* mutants exhibit a diminished DA caliber at 32-72 hpf, which expands less up to 48 hpf and subsequently contracts more by 72 hpf compared to their WT-like sibs (**Figures 1B, F, H**). Finally, the DA caliber fails to widen in *sih* mutants, retaining its 24 hpf size (**Figures 1C, F, I**) in alignment with previous observations[66]. Compared to the *sih* mutant, the residual DA caliber modulation of *plxnd1* nulls is consistent with the idea that other flow-responsive regulators of DA caliber are also active during this period.

To investigate the reasons behind the differences in DA caliber between *plxnd1^fov01b^* and *plxnd1^GAP^*mutants, we compared *plxnd1^Df(Chr08)fs31^* mutants and *plxnd1* morphants at 32 hpf, which share the angiogenic misguidance[58, 59] (**Figures S1A, C-G**) and vascular caliber defects of *plxnd1^fov01b^* fish (**Figure S3**). These observations rule out various potential causes for the reduced DA caliber in *plxnd1^fov01b^*mutants, including impairment of the receptor’s GAP function, misguided angiogenic vessels yielding an abnormal blood flow circuit, linked secondary mutations, or genetic compensation triggered by the mutant transcript’s nonsense-mediated RNA decay[67].

To define the role of blood flow in the *plxnd1*-mediated regulation of DA caliber, we eliminated the circulation from *plxnd1^fov01b^* and *plxnd1^GAP1^* fish using *sih*; *plxnd1^GAP1^* and *sih*; *plxnd1^fov01b^*double mutants. We found that both kinds of double mutants and their single *sih* mutant sibs have identically narrow DA calibers (**Figures 1D-E**), indicating that *plxnd1* regulates the caliber of the DA in a manner dependent on blood circulation.

Lastly, we used a rescue approach to identify which tissue necessitates *plxnd1* to regulate DA caliber. Since *plxnd1* expression includes the endothelium, where it is necessary to guide the vasculature’s anatomical patterning[59, 68-76], and cultured human ECs use *PLXND1* to sense fluid forces[29], we asked if the stable endothelial expression of an HA-tagged form of WT Plxnd1[77] via a *Tg(fliep:2xHA-plxnd1)* transgene suffices to normalize the DA caliber deficit and angiogenic misguidance of *plxnd1^fov01b^* mutants. Indeed, *plxnd1^fov01b^*; *Tg(fliep:2xHA-plxnd1)/+* fish display proper angiogenic guidance and an expanded DA caliber (**Figures 1J-N**). These findings highlight the endothelial necessity for *plxnd1* in regulating both features.

### Plxnd1 promotes the expression of *klf2a* in the DA endothelium in response to blood flow

The *KLF2* (*Krüppel-like factor 2*) genes encode zinc-finger transcription factors expressed in various tissues. Their expression in ECs responds to blood flow, contributing to the atheroprotective adaptation of blood vessels to hemodynamic forces, including maintaining their integrity[30-32, 78-83].

Zebrafish have two *klf2* paralogs, *klf2a* and *klf2b*, with the flow-inducible endothelial expression of *klf2a* being well established [32, 33, 52, 84-89]. To measure the effect of circulatory flow and *plxnd1* activities on *klf2* DA endothelial expression levels with cellular resolution, we used the *klf2a* transcriptional reporter *Tg(klf2a:H2b- EGFP)^ig11^*. In this transgene, a 6 kb *klf2a* promoter fragment drives the expression of a nuclear-targeted histone-EGFP fusion protein[87]. We masked the latter’s endothelial signals with the *Tg(kdrl:nls-mCherry)^y173^*blood endothelium nuclear reporter[90].

We found that the fluorescence levels of *Tg(klf2a:H2b-EGFP)^ig11^* in the ECs of the DA at 84 hpf are similar between *plxnd1^GAP1^* mutants and their WT-like sibs (**Figures 2A-B, F**). Notably, embryos of both genotypes exhibit properly sized DA calibers (**Figures 1A, F, G**). In contrast, both *plxnd1^fov01b^* (**Figures 2C, F**) and *sih* mutants lacking circulation (**Figures 2D, F**) show lower levels of the *klf2a* reporter and reduced DA calibers (**Figures 1B, C, H, I**). In particular, *sih* mutants show the most significant decrease in both aspects. Remarkably, comparing single *sih* mutants and their *plxnd1^fov01b^*; *sih* double mutant siblings reveals similar reductions in the endothelial levels of the *klf2a* reporter (**Figures 2E-F**). Hence, *plxnd1* regulates the expression of endothelial *klf2a* in a manner that relies on blood flow. This finding is consistent with the dampened increase in *Klf2* mRNA levels observed in 2D cultures of murine ECs upon *PLXND1* suppression under athero-protective fluid shear stress[29].

**Figure 2.**
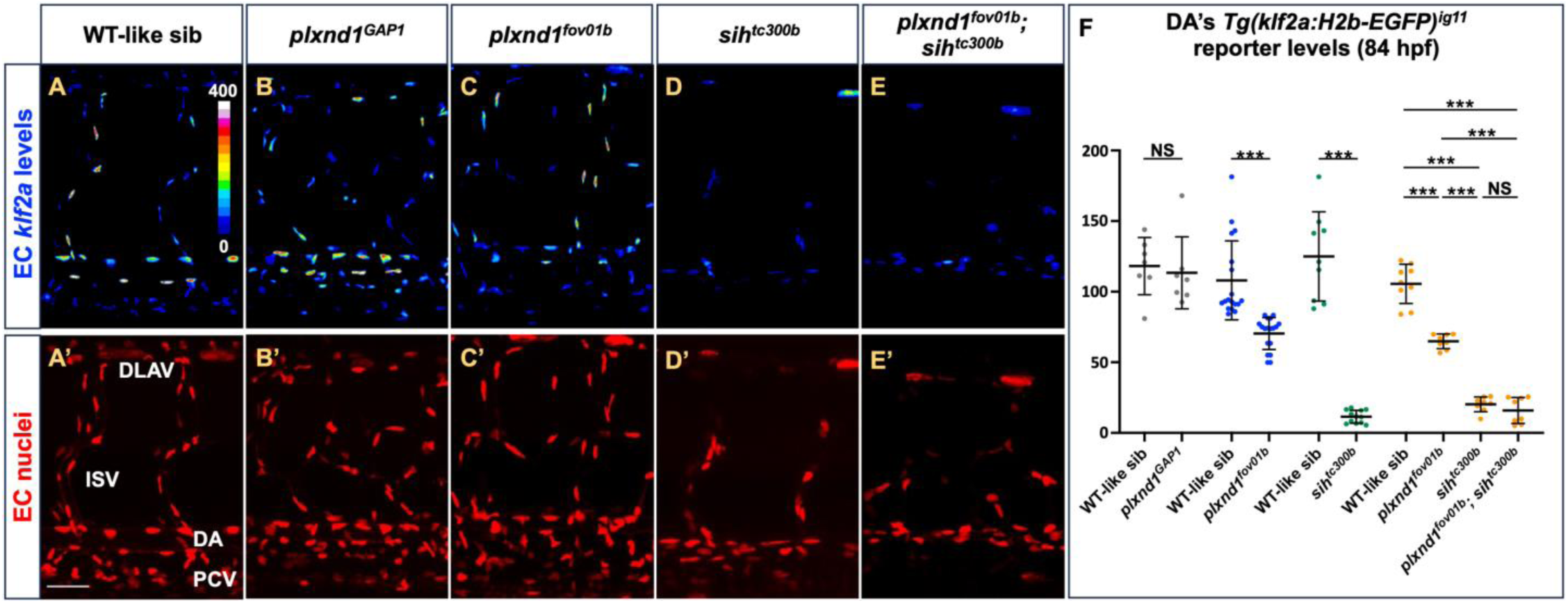
Plxnd1, but not its GAP activity, promotes the DA’s endothelial expression of the *klf2a* transcriptional reporter *Tg(klf2a:H2b-EGFP)^ig11^* in a circulation-dependent manner. (**A-F**) **Data from** live embryos of various genotypes at 84 hpf. (**A-E’**) Lateral views of the trunk. The dorsal side is up, and the anterior side is left. Endothelial nuclear signals from the *Tg(klf2a:H2b- EGFP)^ig11^* reporter scaled by intensity (A, B, C, D, E); intensity scale in (A). Endothelial nuclear signals (red) from the *Tg(kdrl:nls-mCherry)^y173^* reporter, used to mask the *Tg(klf2a: H2b-EGFP)^ig11^* signals with tissue specificity (A’, B’, C’, D’, E’). (**A-A’**) WT sib. The scale bar (horizontal white line) is 40 um (A’). (**B-B’**) *plxnd1^GAP1^* mutant. (**C-C’**) *plxnd1^fov01b^* mutant. (**D-D’**) *sih^tc300b^* mutant. (**E-E’**) *plxnd1^fov01b^; sih^tc300b^* double mutant. (**F**) Graph of live DA’s EC fluorescence intensity of the *Tg(klf2a:H2b-EGFP)^ig11^ reporter.* <u>Statistical measures are</u> <u>as follows</u>. Dots represent individual data points (color-coded to indicate siblings). The long and thick horizontal lines represent the means. The short and thin horizontal lines represent the error bars (SD). Significance levels as NS > 0.05, *p ≤ 0.05, ∗∗p ≤ 0.01, ∗∗∗p ≤ 0.001. Unpaired two-tailed Student’s *t*-test. Comparison of WT-like sibs (n = 7; WTs and *plxnd1^GAP1^*/+) vs. *plxnd1^GAP1^* mutants (n= 7), gray dots, p=0.7045. Comparison of WT-like sibs (n = 18; WTs and *plxnd1^fov01b^*/+) vs. *plxnd1^fov01b^* mutants (n = 18), blue dots, p<0.0001. Comparison of WT-like sibs (n = 9; WTs and *sih^tc300b^*/+) vs. *sih^tc300b^* mutants (n = 12), green dots, p<0.0001. Comparison of WT-like sibs (n = 9; WTs, *plxnd1^fov01b^*/+, *sih^tc300b^*/+, and *plxnd1^fov01b^*/+; *sih^tc300b^*/+)*, plxnd1^fov01b^* mutants (n = 8; *plxnd1^fov01b^* and *plxnd1^fov01b^*; *sih^tc300b^*/+), *sih^tc300b^* mutants (n = 8; *sih^tc300b^* and *plxnd1^fov01b^*/+; *sih^tc300b^*), and *plxnd1^fov01b^*; *sih^tc300b^* double mutants (n = 8), orange dots. WT-like sibs vs. *plxnd1^fov01b^*, p<0.0001. WT-like sibs vs. *sih^tc300b^*, p<0.0001. WT-like sibs vs. *plxnd1^fov01b^*; *sih^tc300b^* double mutants, p<0.0001. *plxnd1^fov01b^* vs. *sih^tc300b^*, p<0.0001. *plxnd1^fov01b^* vs. *plxnd1^fov01b^*; *sih^tc300b^* double mutants, p<0.0001. *sih^tc300b^* vs. *plxnd1^fov01b^*; *sih^tc300b^* double mutants, p=0.2592.

Finally, WT embryos show brighter endothelial *klf2a* reporter fluorescence at the DA’s floor (**Figure 2A**). This observation fits the prediction of higher shear stress in this ventral region arising from this vessel’s dorsal proximity to the rigid notochord and ventral vicinity to the compliant Posterior Cardinal Vein (PCV) and the elastic medium around this vessel[66].

### *klf2* activity is necessary for proper DA caliber

Our data show that Plxnd1 positively affects DA caliber and endothelial *klf2a* expression. These effects depend on blood circulation and can be distinguished from the receptor’s role in guiding blood vessels (**Figures 1-2**), suggesting that Plxnd1 relies on different downstream molecules to execute its distinct vascular roles.

Various observations highlight Klf2 as a candidate Plxnd1 mechano-effector during DA caliber regulation. First, both blood flow (see[32, 33, 52, 84-89]) and *plxnd1* control the endothelial expression of *klf2a* in the DA (**Figure 2**). Second, albeit its transcript’s *in situ* visualization remains unpublished, *klf2b* also expresses endothelially[31, 91, 92], suggesting its flow- and Plxnd1-responsiveness and functional redundancy with its paralog, *klf2a*. Third, *klf2* activity, like circulatory flow but unlike *plxnd1*, is dispensable for guiding the early anatomical patterning of blood vessels[56, 58, 59, 93]. Finally, accumulating evidence implicates this transcription factor in vessel size regulation. Mammalian and piscine studies have found that mutations in Cerebral Cavernous Malformations (CCM) complex components and their partners induce pathogenic blood vessel dilations with elevated *klf2* endothelial expression, and suppressing *klf2* levels ameliorates these vascular lesions[34, 36, 37, 94-96]. Additionally, *klf2a^-^*and *klf2a^-^*; *klf2b^-/+^* fish mutants show axial vessels with progressively smaller outer diameters[35], consistent with the possibility that these animals have reduced vascular calibers.

The hypothesis that Klf2 is a Plxnd1 mechano-effector predicts its role as a positive regulator of the DA caliber. To test this notion, we used the *klf2a* (*klf2a^y616^*)[86] and *klf2b* (*klf2b^sa43252^*)[97] putative loss-of-function alleles. Indeed, our 72 hpf measurements show that the *klf2a^y616^*and *klf2b^sa43252^* single mutants have a reduced DA caliber, with the *klf2* double mutants showing a greater DA caliber deficit (**Figures 3A-E**). Relative to their respective WT-like sibs, the *klf2* double mutants show a greater reduction in DA caliber than *plxnd1^fov01b^* fish (**Figure 3F**), consistent with the notion that the latter exhibits diminished *klf2* expression (**Figures 2C, F**).

**Figure 3.**
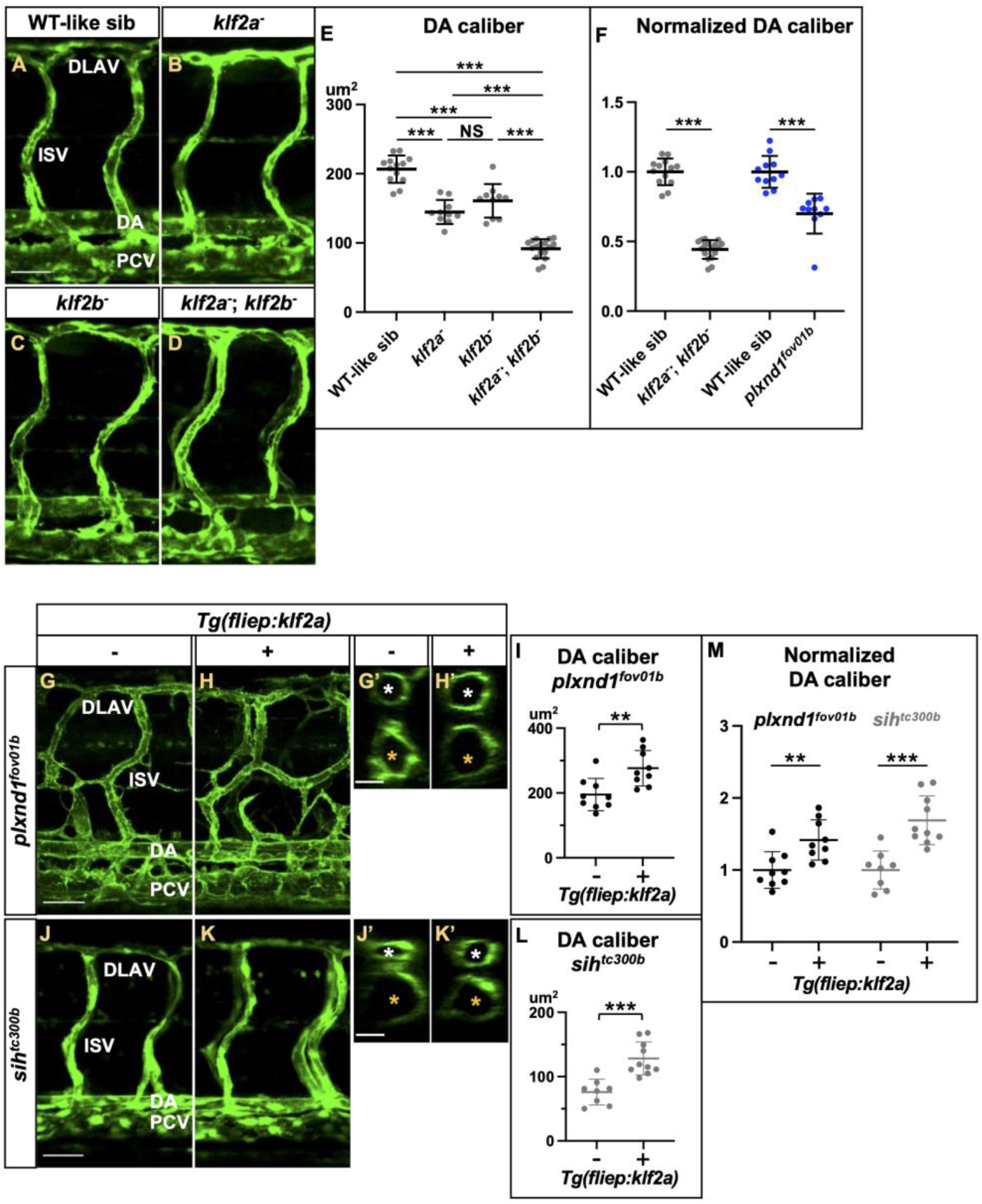
Loss of *klf2* genes reduces the DA caliber, while forced endothelial expression of *klf2a* expands the DA caliber of both *plxnd1^fov01b^* and *sih* mutants without affecting angiogenic patterning. (**A-F**) Images (A-D) and quantifications (E-F) from 72 hpf fixed embryos. (**A-D**) Lateral views of the trunk’s vasculature. The dorsal side is up, and the anterior side is left. Endothelium (green), *Tg(fli1:EGFP)^y1^*. (**A**) WT-like sib. The scale bar (horizontal white line) is 40 μm long. (**B**) *klf2a^y616^* mutant. (**C**) *klf2b^sa43252^* mutant. (**D**) *klf2* double mutant (*klf2a^y616^*; *klf2b^sa43252^*). (**G-M**) Images (G-K’) and quantifications (I, L-M) from 72-76 hpf live embryos. (**G-H’**) Endothelium (green), *Tg(fli1a:lifeactGAP)^mu240^*. (**J-K’**) Endothelium (green), *Tg(fli1:EGFP)^y1^*. (**G-H, J-K**) Lateral views of the trunk’s vasculature. The dorsal side is up, and the anterior side is left. (**G, J**) The scale bar (horizontal white line) is 40 μm long. (**G’, H’, J’, K’**) Representative DA and PCV cross-sections reconstructed from confocal microscopy images. The dorsal side is up, and the left side is left. White asterisks mark the DA. Yellow asterisks mark the PCV. (**G, G’**) *plxnd1^fov01b^* mutant. (**H, H’**) *plxnd1^fov01b^*; *Tg(fliep:klf2a)* mutant. (**J, J’**) *sih^tc300b^* mutant. (**K, K’**) *sih^tc300b^*; *Tg(fliep:klf2a)* mutant. (**G’, J’**) The scale bar (horizontal white line) is 15 μm long. (**E-F, I, L-M**) Quantifications. <u>Statistical measures are as follows</u>. The dots represent individual data points. The horizontal lines represent the means. The error bars denote the SD. Significance levels as *p≤0.05, ∗∗p≤0.01, ∗∗∗p≤0.001, Unpaired two-tailed Student’s *t*-test. (**E**) DA caliber in WT-like sibs (homozygous WTs and heterozygotes for the *klf2a^y616^* or *klf2b^sa43252^* alleles), *klf2a^y616^*and *klf2b^sa43252^* single and double mutants. WT-like sibs (homozygous WTs and heterozygotes for the *klf2a^y616^* or *klf2b^sa43252^* alleles) (n = 13). *klf2a^y616^* (n = 10). *klf2b^sa43252^* (n = 10). *klf2a^y616^* and *klf2b^sa43252^* double mutants (n = 17). WT-like sib vs. *klf2a^y616^*, p<0.0001. WT-like sib vs. *klf2b^sa43252^*, p<0.0001. WT-like sib vs. *klf2a^y616^*; *klf2b^sa43252^*, p<0.0001. *klf2a^y616^* vs. *klf2b^sa43252^*, p=0.9787. *klf2a^y616^* vs. *klf2a^y616^*; *klf2b^sa43252^*, p<0.0001. *klf2b^sa43252^* vs. *klf2a^y616^*; *klf2b^sa43252^*, p<0.0001. (**F**) Normalized DA calibers in the *klf2* double and *plxnd1^fov01b^* single mutants and in the latter’s WT-like sibs (homozygous WTs and *plxnd1^fov01b/+^* heterozygotes). WT-like sib (n = 13) vs. *klf2a^y616^*; *klf2b^sa43252^* (n = 17), p<0.0001. WT-like sib (n = 11) vs. *plxnd1^fov01b^* (n = 10), p<0.0001. (**I**) DA caliber in WT-like sibs (homozygous WTs and *plxnd1^fov01b/+^* heterozygotes) and *plxnd1^fov01b^* mutants with and without the *Tg(fliep:klf2a)* transgene. *plxnd1^fov01b^* (n = 10) vs. *plxnd1^fov01b^*; *Tg(fliep:klf2a)* (n = 10), p=0.0044. (**L**) DA calibers in *sih^tc300b^* mutants with and without the *Tg(fliep:klf2a)* transgene. *sih^tc300b^* (n = 8) vs. *sih^tc300b^*; *Tg(fliep:klf2a)* (n = 10), p=0.0002. (**M**) Normalized DA calibers in *plxnd1^fov01b^* and *sih^tc300b^* mutants and in the WT-like sibs of the former (*plxnd1^fov01b/+^* heterozygotes).

### Forced endothelial expression of *klf2*a normalizes the DA caliber of *plxnd1^fov01b^* and *sih* mutants

If Klf2 is a Plxnd1 mechano-effector during DA caliber regulation, it should function in the same tissue as the receptor and compensate for its absence. To test this hypothesis, we made a *Tg(fliep:klf2a)* transgene to force the expression of Klf2a in ECs. As predicted, this transgene normalizes the DA caliber deficit of *plxnd1^fov01b^* mutants (**Figures 3G’-I**) while leaving their angiogenic patterning defects intact (**Figures 3G-H**).

To investigate whether artificially increasing Klf2 expression in ECs enlarges the DA caliber in a circulation-dependent way, we compared the 72 hpf DA caliber of *sih* mutants and their siblings with and without the *Tg(fliep:klf2a)* transgene. Despite their lack of blood flow, forced endothelial upregulation of *klf2a* levels expands the DA caliber in *sih* mutants (**Figures 3J’-L**), indicating that blood flow promotes the expression, rather than the function, of Klf2 to enact vascular caliber expansion. Increasing the expression of *klf2a* did not alter the anatomical pattern of the angiogenic blood vessels in the trunk of *sih* mutants (**Figures 3J-K**).

Remarkably, while artificially expressing *klf2a* in endothelial cells increases the DA caliber in both *plxnd1^fov01b^* and *sih* mutants to a similar extent (**Figure 3M**), it has no impact on the DA caliber of their WT-like sibs.

### Blood flow, Plxnd1, and Klf2 limit EC abundance, but the DA caliber reduction that occurs in their absence is not due to this vessel’s hyperplasia

Different endothelial cell mechanisms drive flow-dependent vascular caliber adjustments. An increase in either the number of ECs[88, 98-105] or their area[52, 104, 106, 107] can expand vascular caliber. Conversely, increased EC elongation and alignment to the vessel’s axis can narrow its lumen[52].

To understand how Plxnd1 increases DA caliber in response to circulation at the cellular level, we first manipulated *plxnd1* activities and blood flow. To achieve this goal, we assessed the DA’s EC abundance in *plxnd1^GAP1^*, *plxnd1^fov01b^*, and *sih* mutants and their WT-like sibs, scoring the embryos before (24 hpf) and after (32, 48, and 84 hpf) the circulation’s establishment. We used the *Tg(fli:nEGFP)^y7^*pan-endothelial nuclear reporter[100] to quantify EC nuclei within a specified DA segment, posteriorly delimited by the yolk’s extension end. We defined the count as DA’s ECs per somite (DA’s ECs/somite).

We noticed a gradual decrease in EC abundance among the WT-like sibs, indicating that the DA caliber expansion between 24-48 hpf typically happens without increasing the number of DA’s ECs (**Figures S4A-C, F**). EC abundance also diminishes in both *plxnd1* mutants, but more slowly (**Figures S4A-B, G-H**). In contrast, EC abundance increases gradually in the DA of *sih* mutants (**Figures S4C, I**). The timing, direction, and magnitude of EC abundance change in these three mutants (**Figures S4A-C**), alongside their distinct patterns of DA caliber evolution (**Figures 1A-C**), argues that the narrow DA caliber of *plxnd1^fov01b^* and *sih* mutants is not due to this vessel’s hyperplasia (the two sections below provide additional support for this notion).

Finally, to determine whether the loss of *klf2* leads increases the DA’s EC abundance, as observed upon *plxnd1* inactivation, we compared single (*klf2a*^-^ and *klf2b^-^*) and double *klf2* mutants at 84 hpf. The single and double *klf2* mutants display elevated EC abundance, with the latter genotype showing a significantly stronger phenotype. However, EC numbers are higher in *plxnd1^fov01b^*fish (**Figures S4D-E, F, H, J-L**).

### Diminished circulatory shear stress limits the DA’s caliber expansion without affecting EC abundance

To further clarify how circulatory forces impact the DA’s caliber and EC abundance, we evaluated the impact of lessening circulatory fluid shear stress (by reducing blood viscosity) on these two features over time and its effect on heartbeat frequency (**Figure S5**). Specifically, we lowered the hematocrit by impairing the function of the *gata1a* gene, a critical transcriptional driver of primitive erythrocyte differentiation. Homozygous *gata1a^m651^*mutants (previously known as *vlad tepes* or *vlt*) lack almost all red blood cells (RBCs)[108] and die between 8- 15 dpf[109]. Notably, at 48 hpf (when the DA caliber is the widest in WTs), *gata1a^m651^* fish have a beating heart with normal morphology, including an atrioventricular canal (AVC) with the proper number of endocardial cells[110], suggesting that the mutant’s cardiac function is grossly unaffected by 48 hpf.

Consistent with this notion, we found that *gata1a^m651^* mutants and their WT-like siblings have identical heartbeat frequencies at 50 hpf (**Figures S5C**). Both genotypes display similar DA calibers at 24 hpf before the heartbeat’s onset (**Figure S5A**), indicating that the intravascular entry of RBCs ongoing at this stage is dispensable for DA lumenization and sizing; see[111]. However, in the presence of circulation, the DA of *gata1a^m651^* fish expands significantly less at 32 and 48 hpf and fails to narrow by 72 hpf. Hence, in *gata1a^m651^* mutants, the DA caliber is constant from 32-72 hpf. Despite these mutants’ DA caliber deficit, their EC abundance is indistinguishable from their WT-like siblings at 24-48 hpf. However, *gata1a^m651^* mutants display a mild increase in EC abundance by 84 hpf (**Figure S5B**). These findings argue that the *gata1a^m651^* mutants’ reduced circulatory shear stress limits the circulation-induced expansion phase of the DA that culminates at 48 hpf in WTs. Importantly, this deficit is not due to changes in EC abundance. Despite robust blood flow, we note that *plxnd1^fov01b^* mutants also exhibit limited DA caliber expansion at 32 and 48 hpf.

### Plxnd1 increases the DA caliber by promoting the enlargement of ECs through Klf2

Our research suggests that Plxnd1 responds to circulatory forces and, via KLF2, influences the size, shape, or alignment of ECs, ultimately controlling the DA’s caliber. To measure these parameters and differentiate between potential cellular mechanisms, we created the *Vessel Analyzer* app. Using images of the endothelial membrane and a junctional marker, the software performs semi-automated single EC morphometry analysis, extracting data on cell length, width, area, and volume, their 2D and 3D shape (elongation and anisotropy of the inertia moment, respectively), and their alignment to the vessel’s axis. Please see the explanatory diagrams of these features in **Figure 4A**.

**Figure 4.**
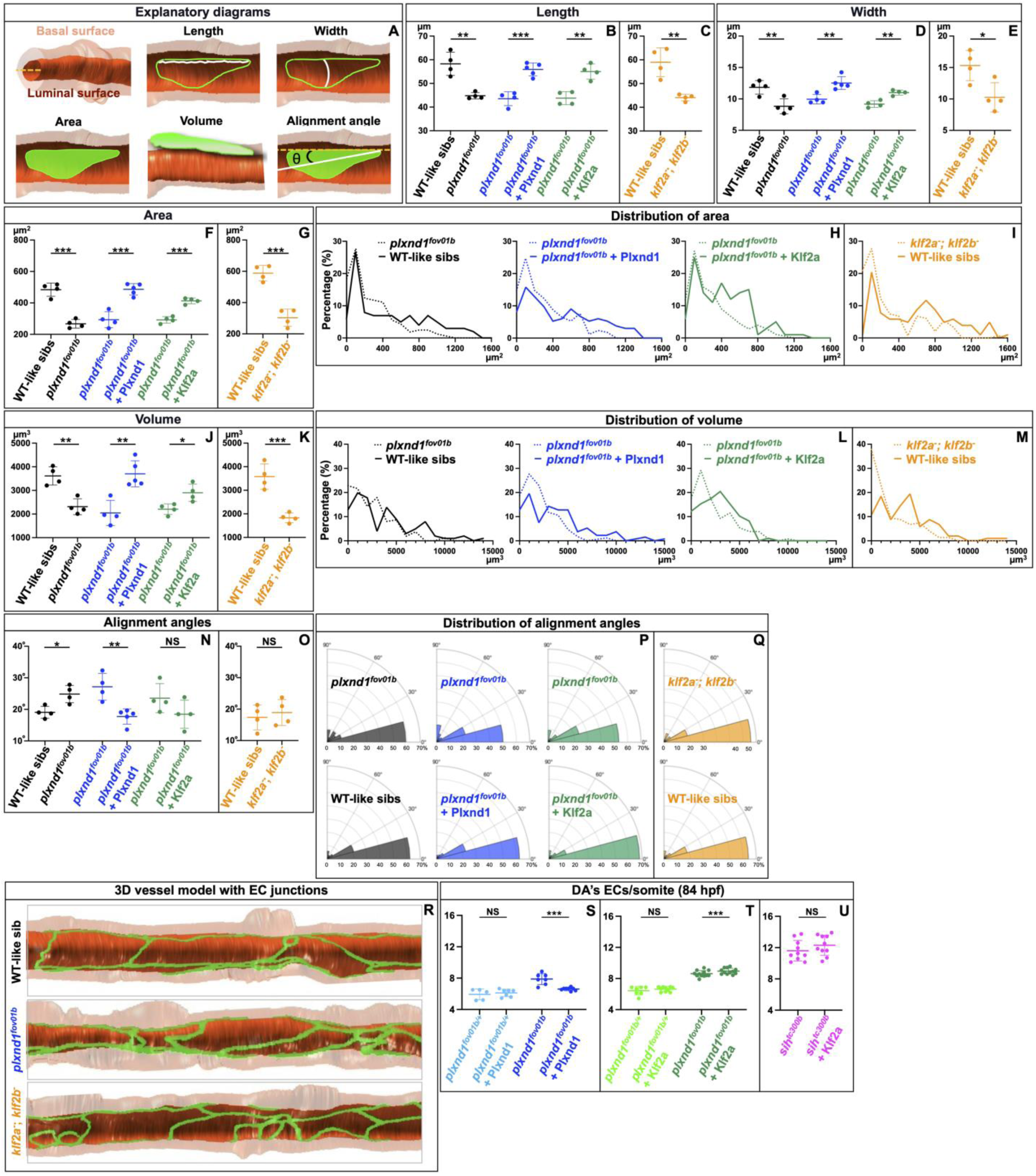
Plxnd1 expands the DA caliber by enlarging the area of ECs via Klf2. (**A**) Explanatory diagrams. On the top left is the <u>location of basal and luminal surfaces in the DA</u>, with the vessel’s longitudinal axis (yellow dashed line) projecting out the page at an angle. The other illustrations define EC morphometric parameters within a DA oriented horizontally. <u>Length</u> (white line): distance along the vessel’s lumen connecting the two points within the cell’s perimeter (shown in green) furthest apart along the vessel’s axis. <u>Width</u> (white line): the transversal length connecting the two most distant points within the cell’s perimeter (outlined in green) along the vessel’s lumen. <u>Area</u> (green surface): measured at the vessel’s luminal side. <u>Volume</u> (green form with shadowing): calculated by integrating the cell’s height between the luminal and basal areas (to ease comprehension, the DA has been rotated 90° around the vessel’s longitudinal axis compared to the other illustrations of EC morphometry). <u>Alignment angle or angle with the vessel’s axis (θ</u>): measured as the angle between the cell’s minor principal axis of rotation (black continuous line) and the vessel’s longitudinal axis (yellow dashed line). See **METHOD DETAILS** (**Zebrafish: imaging and quantification of EC morphometry with the *Vessel Analyzer* app**). <u>Comparisons</u> <u>of EC morphometry in live embryos</u>. (**B, D, F, H, J, L, N, P**) Black: WT-like sibs vs. *plxnd1^fov01b^*. Blue: *plxnd1^fov01b^* vs. *plxnd1^fov01b^* + Plxnd1 (*Tg(fliep:2xHA-plxnd1)*). Green: *plxnd1^fov01b^* vs. *plxnd1^fov01b^* + Klf2a (*Tg(fliep:klf2a)*). (**C, E, G, I, K, M, O, Q**) Orange: WT-like sibs vs. *klf2a^-^*; *klf2b^-^*, p=0.0031. (**B-C**) Graphs of mean EC length values. (B) Comparison of WT-like sibs vs. *plxnd1^fov01b^*, p=0.0018. *plxnd1^fov01b^* vs. *plxnd1^fov01b^* + Plxnd1 (*Tg(fliep:2xHA-plxnd1)*), p=0.0003. *plxnd1^fov01b^* vs. *plxnd1^fov01b^* + Klf2a (*Tg(fliep:klf2a)*), p=0.0018. (C) WT-like sibs vs. *klf2a^-^*; *klf2b^-^*, p=0.0031. (**D-E**) Graphs of mean EC width values. (D) Comparison of WT-like sibs vs. *plxnd1^fov01b^*, p=0.0094. *plxnd1^fov01b^* vs. *plxnd1^fov01b^* + Plxnd1 *Tg(fliep:2xHA-plxnd1)*), p=0.0041. *plxnd1^fov01b^* vs. *plxnd1^fov01b^* + Klf2a (*Tg(fliep:klf2a)*), p=0.0016. (E) WT-like sibs vs. *klf2a^-^*; *klf2b^-^*, p=0.0231. (**F-G**) Graphs of mean EC area values. (F) Comparison of WT-like sibs vs. *plxnd1^fov01b^*, p=0.0001. *plxnd1^fov01b^* vs. *plxnd1^fov01b^* + Plxnd1 (*Tg(fliep:2xHA-plxnd1)*), p=0.0003. *plxnd1^fov01b^* vs. *plxnd1^fov01b^* + Klf2a (*Tg(fliep:klf2a)*, p=0.0001. (G) WT-like sibs vs. *klf2a^-^*; *klf2b^-^*, p=0.0003. (**H-I**) Histograms of EC area distribution. X axis, EC area. Y axis, percentage of ECs scored. (**J- K**) Graphs of mean EC volume values. (J) Comparison of WT-like sibs vs. *plxnd1^fov01b^*, p=0.0022. *plxnd1^fov01b^* vs. *plxnd1^fov01b^* + Plxnd1 (*Tg(fliep:2xHA-plxnd1)*), p=0.0026. *plxnd1^fov01b^* vs. *plxnd1^fov01b^* + Klf2a (*Tg(fliep:klf2a)*), p=0.0180. (K) WT-like sibs vs. *klf2a^-^*; *klf2b^-^*, p=0.0009. (**L-M**) Histogram of EC volume distribution. X axis, EC volume. Y axis, percentage of the ECs scored. (**N-O**) Graphs of the mean EC alignment angles. (N) Comparison of WT-like sibs vs. *plxnd1^fov01b^*, p=0.0128. *plxnd1^fov01b^* vs. *plxnd1^fov01b^* + Plxnd1 (*Tg(fliep:2xHA-plxnd1)*), p=0.0039. *plxnd1^fov01b^* vs. *plxnd1^fov01b^* + Klf2a (*Tg(fliep:klf2a)*), p=0.1613. (O) WT-like sibs vs. *klf2a^-^*; *klf2b^-^*, p=0.6077. (**P-Q**) Quarter-polar histogram plots of EC alignment angle distribution (binned in 15° intervals). R axis, percentage of scored ECs. Theta axis, 0-90°. (**B-Q**) For all genotypes, n = 4, except for *plxnd1^fov01b^* + Plxnd1 (*Tg(fliep:2xHA-plxnd1)*), in which n = 5. (**R**) 3D vessel models with EC junctions made by the *Vessel Analyzer App*. Top panel, WT-like sib. Middle panel, *plxnd1^fov01^*. Bottom panel, *klf2* (*klf2a^-^*; *klf2b^-^*) double mutants. (**S-U**) Graphs comparing the DA’s EC abundance per somite at 84 hpf in sibling embryos. (S) *plxnd1^fov01b/+^* (n = 5) vs. *plxnd1^fov01b/+^* + Plxnd1 (*Tg(fliep:2xHA-plxnd1)*), (n = 7); p=0.3387. *plxnd1^fov01b^* (n = 7) vs. *plxnd1^fov01b^* + Plxnd1 (*Tg(fliep:2xHA- plxnd1)*), (n = 8); p=0.0003. (T) *plxnd1^fov01b/+^* (n = 7) vs. *plxnd1^fov01b/+^* + Klf2a (*Tg(fliep:klf2a)*), (n = 10); p=0.3387. *plxnd1^fov01b^* (n = 7) vs. *plxnd1^fov01b^* + Klf2a (*Tg(fliep:klf2a)*), (n = 10); p=0.0503. (U) *sih^tc300b^* (n = 10) vs. *sih^tc300b^*+ Klf2a (*Tg(fliep:klf2a)*), (n = 10); p=0.2429. <u>Statistical measures are as follows</u>. (**B-G**, **J-K**, **N-O**) Dots denote means from individual larvae. (**S-U**) Dots denote the average EC abundance values of individual larvae. (**B-G**, **J-K**, **N-O**, **S-U**) Long and thick horizontal lines represent the means. Short and thin horizontal lines, error bars (SD). Significance levels as *p≤0.05, ∗∗p≤0.01, ∗∗∗p≤0.001), unpaired two-tailed Student’s *t*-test.

We used the *TgKI(tjp1a-eGFP)^pd1252^*knock-in line as the junctional marker for these cellular morphometry analyses. This reporter produces a functional fluorescent Tjp1a fusion protein from the *tjp1a* locus, a *zonula occludens-1* (*zo-1*) ortholog[112]. This gene encodes a bicellular tight junction component of many cell types, including the endothelium[112-116]. To select the junctions in the latter tissue, we masked the *TgKI(tjp1a- eGFP)^pd1252^* signal with the *Tg(kdrl:mRFP-F)^y286^*endothelial membrane marker[117] (**Figures S6A-A”**).

Our analyses of live *plxnd1*, *klf2*, and *sih* mutants and their WT-like sibs at 84 hpf revealed the following. EC morphometry is unaffected in the adequately sized DA of *plxnd1^GAP1^* fish, except for a mild, albeit statistically significant, change in cell area (a 15.6% reduction, **Figures S7A-E** and **Table S2**). In contrast, the ECs lining the narrow DA of *plxnd1^fov01b^*and *klf2* double mutants share considerable and statistically significant reductions in the mean values for length (∼23% and ∼25%, **Figures 4B-C**, **S7A**, and **Table S2**), width (∼25% and ∼33%, **Figures 4D-E, S7B,** and **Table S2**), area (∼55% and ∼48%, **Figures 4F-G**, **S7C,** and **Table S2**), and volume (∼36% and ∼49%, **Figures 4J-K**, **S7D,** and **Table S2**). However, neither mutant exhibits statistically significant changes in EC shape (**Table S2**). The dispensability of piscine *plxnd1* for EC elongation in response to flow contrasts with findings in 2D cultures of bovine ECs and adult murine aorta in which this morphometric change is receptor-dependent[29]. While this observation might reflect species-specific differences, variables like 2D or 3D endothelial organization, shear stress magnitudes, and exposure periods provide a more likely explanation for this discrepancy.

Notably, the main difference in EC morphometry between *plxnd1^fov01b^* mutants and *klf2* nulls is the former’s EC alignment impairment (from 19.05° in the WT-like sibs to 24.85° in *plxnd1^fov01b^* fish, a ∼30% difference, **Figures 4N-Q**, **S7E,** and **Table S2**). This phenotype fits the alignment defect observed in the aorta of mice with adult endothelial depletion of *Plxnd1* and cultured Bovine Aortic ECs exposed to laminar fluid shear stress with diminished receptor expression[29]. However, unlike *klf2* null fish, *KLF2* knockdown compromises flow-induced alignment in 2D cultures of human ECs[118], perhaps due to differences between species or, more likely, contrasting experimental conditions.

In the endothelium of the narrow DA of *sih* mutants, the Tjp1a-eGFP fusion protein forms discontinuous junctions with variable thickness and accumulates in large puncta (**Figure S6**). This phenotype is consistent with the notion that circulatory forces promote the maturation of endothelial junctions since similar abnormalities occur in the endothelial adherens junctions of *sih* morphants, as revealed by Vascular Endothelial (VE)-cadherin[119]. While the junctional defects of *sih* mutants prevent the evaluation of EC morphometry, the expectation is that their DA supernumerary ECs are also abnormally small.

Taking into account the unusual cellular morphometry observed in the endothelium of *plxnd1^fov01b^*and *klf2* (*klf2a^-^; klf2b^-^*) null mutants and the capacity of the *Tg(fliep:2xHA-plxnd1)* and *Tg(fliep:klf2a)* transgenes to restore the reduced DA caliber in both *plxnd1^fov01b^* and *sih* mutants (**Figures 1B-C, H-I, K, M-N**, and **Figures 3G’-H’, J’- K’, I-M**), we investigated the cellular effects on the DA’s endothelium of forced Plxnd1 and Klf2a expression in *plxnd1^fov01b^* and *sih* mutants at 84 hpf.

In *plxnd1^fov01b^* fish, exogenous expression of Plxnd1 and Klf2a increased the mean values for the ECs’ length (28.4% and 22.8%, **Figures 4B, S7F, K** and **Table S2**), width (25.8% and 17.5%; **Figures 4D, S7G, L** and **Table S2**), area (66% and 36%, **Figures 4F, S7H, M** and **Table S2**), and volume (81% and 18.7%, **Figures 4J**, **S7I, N** and **Table S2**). The data distribution for area and volume also indicates an improvement, with more cells adopting medium to high values and fewer acquiring smaller ones (**Figures 4H-I, L-M**). Neither transgene had statistically significant effects on EC elongation, with augmentations of 0.56% and 3.56% (**Table S2**).

Providing exogenous Plxnd1 reduced the EC’s alignment angle (from 27.14° to 17.72°, a ∼30% difference; see **Figures 4N, P**, **S7J**, and **Table S2**). The *Tg(fliep:2xHA-plxnd1)* transgene also had a minor but statistically significant effect on the ECs’ 3D shape (a 1.3% anisotropy decrease, see **Table S2**). On the other hand, *klf2a* expression improved EC alignment (**Figures 4N, P**, **S7O**, and **Table S2**, a switch from 23.55° to 18.45°) without reaching statistical significance.

To assess the effectiveness of the two transgenes in rescuing the defective EC morphometry of *plxnd1^fov01b^*embryos, we used the mean values of WT-like siblings from a cross of *plxnd1^fov01b/+^* heterozygotes (black label in **Figures 4B, D, F, H, J, L, N, P**) as the benchmark. These assessments suggest that inducing *plxnd1* expression restores normal EC morphometry in *plxnd1^fov01b^* fish more effectively than introducing exogenous *klf2a* (dark blue and dark green labels in **Figures 4B, D, F, H, J, L, N, P**). Specifically, the *Tg(fliep:2xHA-plxnd1)* transgene reinstated all five cellular morphometric characteristics affected in *plxnd1^fov01b^* mutants to WT-like values, namely the length (∼100%), width (∼96%), area (∼95%), volume (∼98%), and alignment (∼93%). However, *Tg(fliep:klf2a)* expression only rescued the four EC morphometry defects shared by *plxnd1^fov01b^* and *klf2* nulls, namely length (∼94%), width (∼92%), area (∼85%), and volume (76%). **Figure 4R** shows *Vessel Analyzer*-generated 3D models of DA segments from WT-like sibs, *plxnd1^fov01b^* fish, and *klf2* (*klf2a^-^; klf2b^-^*) double mutants, with the EC junctions projected onto the vessel’s luminal surface to illustrate some of the effects of *plxnd1* and *klf2* on EC morphometry.

Significantly, introducing exogenous Plxnd1 reduced the excess ECs in the DA of *plxnd1^fov01b^* nulls (**Figure 4S**). In contrast, elevating endothelial Klf2a levels did not alleviate the cellular overabundance in *plxnd1^fov01b^*(**Figure 4T**). and *sih* fish (**Figure 4U**). Nevertheless, forced endothelial expression of Plxnd1 in *plxnd1^fov01b^* embryos (**Figures 1M-N**) and of Klf2a in *plxnd1^fov01b^*(**Figures 3H’, I, M**) and *sih* mutants (**Figures K’, L, M**) expanded their abnormally small DA.

These findings again illustrate that the quantity of ECs in the DA and its caliber can change independently. Moreover, they establish that both Plxnd1 and Klf2 control the DA caliber primarily by influencing the size of ECs, not their quantity. Finally, our data also indicates that Plxnd1, but not its guidance activity, is necessary for optimal EC alignment.

### Engineered microvascular networks (MVNs) made from human ECs respond to circulatory flow by upregulating a reporter of *KLF2* expression, expanding their caliber, and increasing EC size

To determine the evolutionary conservation of the effects of circulatory forces on *KLF2* expression activity, vascular caliber, and EC sizing and bypass the potential confounding effects of developmental changes in cardiac function[44-51] and vascular mural cell investment[65, 86, 120-122], we employed an *in vitro* model.

Specifically, we used ECs carrying a flow-inducible reporter of *KLF2* transcription[123] to construct three-dimensional human MVNs within microfluidic devices and applied flow (or not) using an on-chip microfluidic pump, as in[124] (see also[125, 126]).

To determine the width of vessel lumens, we highlighted the latter by perfusing fluorescent dextran through MVNs. Static MVNs had smaller vessel calibers (54.5 ± 2.7 mm, n=6 MVNs) than MVNs cultured under flow (70.2 ± 11.5 mm, n=5 MVNs), see **Figures 5A-C**. We found that while MVNs cultured under static (no flow) conditions show minimal expression of KLF2-GFP, MVNs exposed to flow express KLF2-GFP robustly (**Figures 5D-F**). Notably, as shown in **Figures 5G-I**, ECs in static MVN have smaller areas (700 ± 132 mm^2^, n=651 cells in 3 MVNs) than ECs in MVNs cultured under flow (1,344 ± 397 mm^2^, n=304 cells in 3 MVNs).

**Figure 5.**
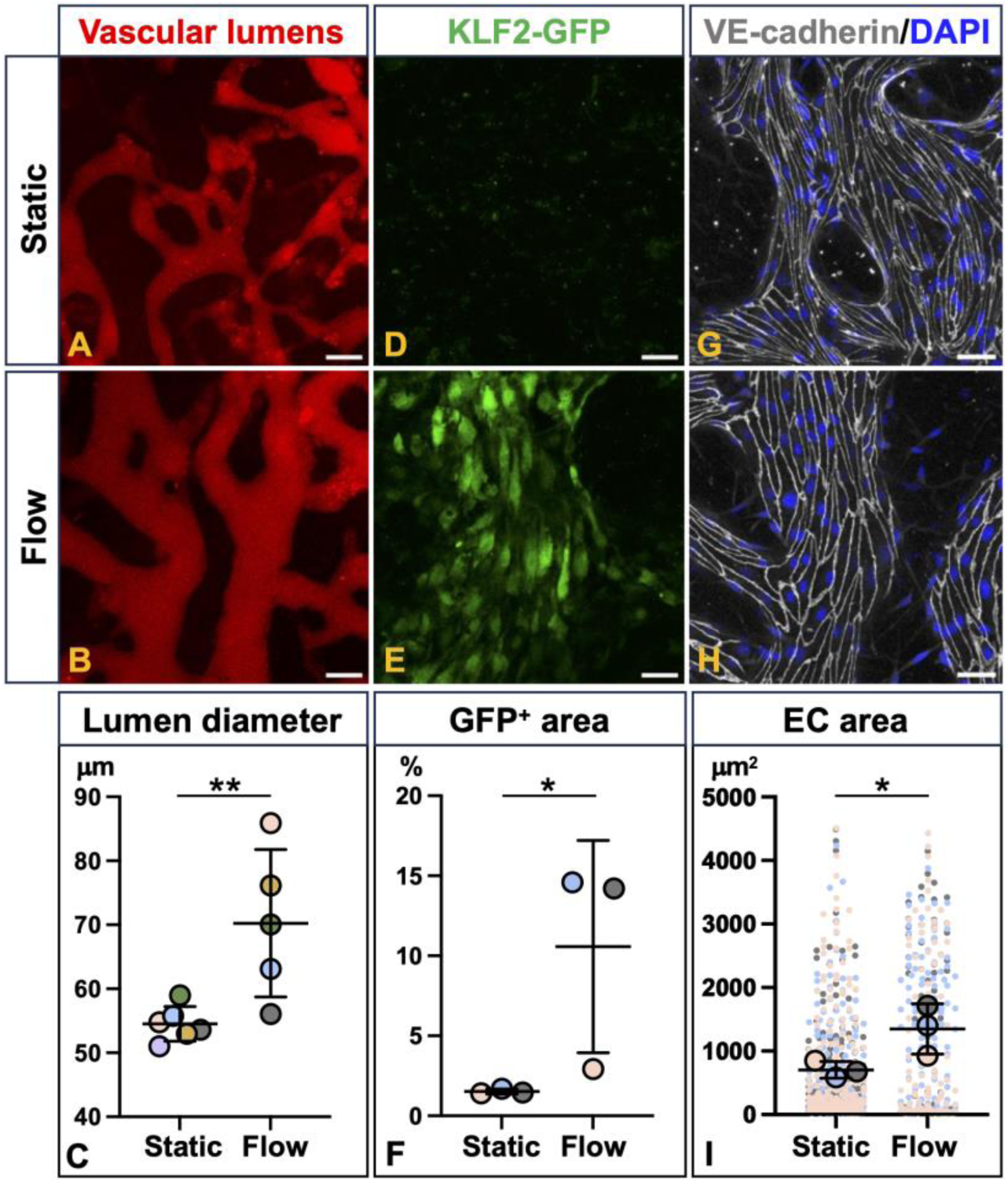
Circulatory flow promotes the activation of a reporter of *KLF2* expression, caliber expansion of vessels, and EC enlargement in MVNs. Representative confocal images (maximum intensity projections) of vascular lumens illuminated with fluorescent dextran. (**A-B**; red, scale bar is 100 µm), KLF2-GFP expression (**D-E**; green, scale bar is 50 µm), and vascular endothelial-cadherin immunofluorescence (VE-cadherin, gray) and DAPI staining (blue) outlining EC perimeters and marking cell nuclei (**G-H**; scale bar is 50 µm) in MVNs under static (**A, D, G**) and flow (**B, E, H**) conditions. (**C**) Graph comparing the average luminal diameter of MVNs cultured under static and flow conditions. p=0.0097 for flow (5 MVNs) vs. static (6 MVNs). (**F**) Graph comparing KLF2-GFP fluorescence levels, expressed as the percentage of GFP-positive imaging area of MVNs cultured under static and flow conditions. p=0.0386 for flow (3 MVNs) vs. static (3 MVNS) conditions. (**I**) Superplot graph[186] compares individual ECs’ areas in MVNs cultured under static and flow conditions. p=0.0280 for flow (3 MVNs, 651 cells) vs. static (3 MVNs, 304 cells) conditions. <u>Statistical measures are as follows</u>. Larger circles denote means (color-coded per MVN). Small dots represent individual EC measurements (color-coded per MVN). Long horizontal lines represent the means of each MVN. Short horizontal lines, error bars (SD). Significance levels as *p≤0.05, ∗∗p≤0.01, ∗∗∗p≤0.001, unpaired one-tailed Student’s *t*-test.

The differential effects of static and flow conditions in the MVNs parallel those observed in the piscine DA of *sih* mutants and WTs, arguing that circulatory forces modulate vascular caliber via similar molecular and cellular effects from fish to humans.

### Murine embryos with endothelial *Plxnd1* deficiency display decreased DA caliber

To investigate whether PLXND1 plays a similar role in regulating the size of the DA in fish and mammals, we selectively deleted *Plxnd1* in the endothelium of mice embryos using the *Tie2^Cre^* deleter line[127] and the floxed *Plxnd1* (*Plxnd1*^flox^) allele[71, 74]. Specifically, we harvested control and *Tie2^Cre/+^;Plxnd1^flox/flox^* embryos from timed matings and made paraffin sections from them, which we processed as follows. We performed Hematoxylin and eosin (H&E) staining (as in[128-130]) or double immunofluorescence for Platelet endothelial cell adhesion molecule (Pecam1 or CD31) and ETS-Related Gene 1 (ERG1) to visualize ECs and their nuclei.

We measured the DA at the cardiac level in cross-sections of E11.25 embryos, when blood flow through the DA is present already[131-135] and*Tie2^Cre/+^;Plxnd1^flox/flox^* embryos exhibit minimal morphological and structural changes in the outflow tract compared to the controls; see[69, 71]. Compared to the control group, we observed a marked reduction in the luminal area and luminal perimeter of the DA in *Tie2^Cre/+^;Plxnd1^flox/flox^*(*Plxnd1^ECKO^*) embryos (**Figure 6**).

**Figure 6.**
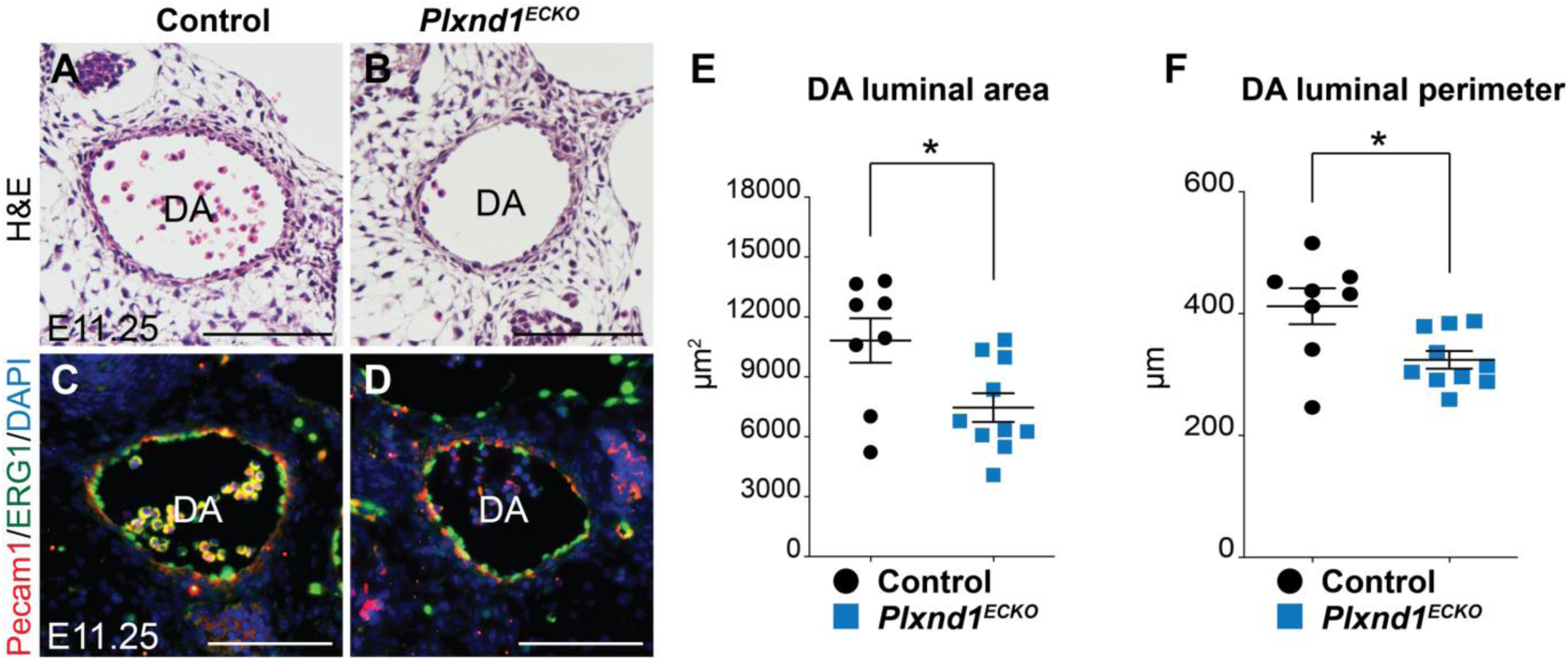
Endothelial *Plxnd1* is necessary for proper DA caliber in murine embryos. (**A-B**) Hematoxylin and eosin (H&E) stained paraffin sections from the cardiac level, showing the DA of E11.25 control (A) and *Plxnd1^ECKO^* (B) embryos. (**C-D**) Cardiac level sections (scale bars, 100 μm) of E11.25 control (C) and *Plxnd1^ECKO^* (D) embryos immunofluorescently stained to visualize the DA’s expression of Pecam1 (red) and ERG1 (green) and counterstained with the DNA stain DAPI (blue) to highlight nuclei. (**A-D**) Scale bars, 100 μm. (**E-F**) Graphs of the DA’s luminal area (E) and luminal perimeter (F) in H&E stained control (n=4, black) and *Plxnd1^ECKO^* embryo sections (n=5, blue). We measured both the right and the left DA of each embryo. <u>Statistical measures are as follows</u>. We report the values as means ± SEM (standard error of the mean). Significant differences, \**p*<0.05; unpaired two-tailed Student’s *t*-test. The two genotypes show statistically significant differences in both area (*p*=0.0186) and perimeter (*p*=0.0111).

Together with our piscine data, these murine findings support the notion that, across vertebrates, *PLXND1* is a positive regulator of the DA’s caliber. Furthermore, in light of the results from our MVN experiments, these observations suggest that the receptor exerts its mechanosensing-dependent effects on vascular caliber regulation using evolutionarily conserved molecular and cellular mechanisms.

## DISCUSSION

Our piscine results indicate that the novel mechanosensory function of Plxnd1 is endothelially required to widen the DA’s luminal opening in response to increased circulatory friction against the vascular wall. We also uncover that the flow-responsive transcription factor KLF2 acts as a paramount mechanosensitive effector of Plxnd1 that enlarges ECs to widen the vessel (**Figure 7**). Furthermore, our data from WT and mutant fish embryos implies that circulatory forces (absent and reduced in *sih* and *gata1a^m651^* mutants, respectively) and their adequate sensing and transduction (impaired in *plxnd1^fov01b^* and *klf2* mutants) are critical for correctly sizing this vessel.

**Figure 7.**
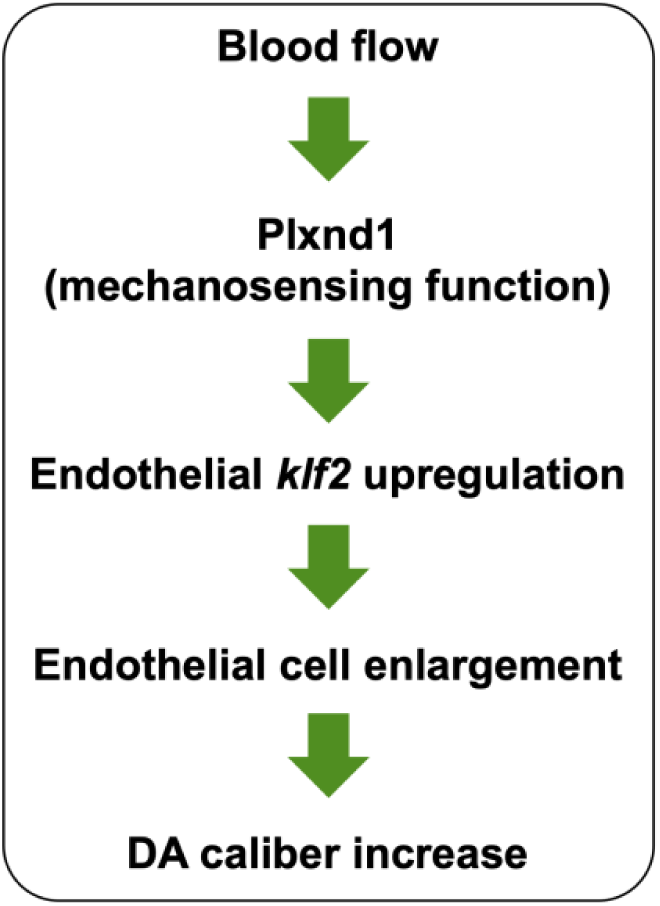
Model of circulation-dependent DA caliber regulation by Plxnd1 via Klf2.

Specifically, we uncovered allele-specific effects of *plxnd1* by comparing the DA calibers of mutants with similar angiogenic misguidance phenotypes, namely *plxnd1^fov01b^*, and *plxnd1^GAP1^*. Before the onset of blood flow, these mutants and their WT-sibs show equivalent DA sizes. Later, however, when circulation is present, these mutants differ, showing reduced and typical DA calibers, respectively. Consistent with the notion that Plxnd1 regulates vascular caliber with flow-dependency, we found that *sih* fish and both double *plxnd1;sih* mutants display equivalent DA reduction phenotypes. The smaller caliber of the DA in circulation-deficient *sih* fish compared to *plxnd1* nulls during this vessel’s expansion phase, culminating at 48 hpf, highlights the involvement of additional flow-dependent regulators of vascular caliber.

These findings also reveal that the role of Plxnd1 in DA caliber regulation is distinct from its vascular guidance function, as the former is unaffected by mutations predicted to block its GAP activity, critical for repulsive signaling[63, 64, 136, 137]. Hence, the narrow DA caliber of *plxnd1^fov01b^* nulls is not secondary to their abnormal circulatory topology, demonstrating *in vivo* that the receptor’s two primary functions are genetically separable. This conclusion aligns with the *in vitro* observation that a Plexin-D1 receptor with the ectodomain locked in the closed conformation lacks mechanosensory activity but still triggers cell collapse in response to the repulsive guidance cue Sema3E[29].

Consistent with the expected requirement for Plxnd1 in the endothelium for angiogenic guidance[58, 59, 69- 72] and the sensing of circulatory forces[29] that modulate DA sizing, we found that forced expression of the WT receptor in this tissue suffices for restoring angiogenic pathfinding and normalizing the reduced DA caliber of *plxnd1^fov01b^* nulls.

Our piscine data also establishes that *plxnd1* is necessary for the efficient circulation-dependent transcriptional upregulation of *klf2a*, a piscine ortholog of human *KLF2*, in the DA’s ECs; see[32, 33, 52, 84-89]. It is likely that the receptor similarly fosters the endothelial expression of *klf2b* and *klf4*[31, 91, 92], given the reduction of *KLF2* and *KLF4* levels in cultured ECs under laminar fluid shear stress upon *PLXND1* attenuation[29]. How the receptor regulates the expression of these transcription factors is unclear but likely involves indirect mechanisms, given its localization to the cell membrane rather than the nucleus[68, 138-141] (but see[142]). Some potential avenues include MAP3K/MEK5/ERK5 pathway activation[143] and inhibition of CCM (Cerebral Cavernous Malformation) proteins[144], p53[145], and p66shc[146], which promote and antagonize these transcription factors endothelial expression.

Additionally, our findings establish a direct connection between *klf2* expression and DA caliber sizing, position Klf2 as a Plxnd1 mechano-effector, and reveal that this transcription factor’s activity is independent of blood flow. First, *klf2a* expression is unaffected in *plxnd1^GAP1^*fish with an adequately sized DA lumen, but it is lower in *plxnd1^fov01^* null mutants with a reduced DA caliber. This observation implies that other flow-responsive regulators drive residual endothelial *klf2a* expression without Plxnd1. Second, *sih* mutants also exhibit reduced *klf2a* levels and decreased DA calibers, and these defects are more robust than those of *plxnd1^fov01^*nulls. Notably, the reduction in *klf2a* levels and DA caliber is similarly substantial in *sih* and double *plxnd1^fov01b^; sih* mutants, indicating that Plxnd1 regulates both features in a circulation-dependent way. Third, single *klf2a* and *klf2b* mutants also have a smaller-than-normal DA caliber, which reduces further in *klf2* double mutants, fitting with these genes’ expected overlapping functions and the reciprocal paralog’s upregulation in the single mutants[93]. Finally, forced endothelial expression of *klf2a* normalizes the DA caliber of both *sih* fish devoid of blood flow and *plxnd1^fov01b^* mutants with circulation. However, exogenous *klf2a* expression does not rescue the *plxnd1^fov01b^* mutants’ angiogenic misguidance, consistent with the presence of adequately navigating angiogenic vessels in *sih* and *klf2* mutants.

Surprisingly, forced endothelial expression of *klf2a* in WT-like embryos does not affect the DA’s caliber. Perhaps the *Tg(fliep:klf2a)* transgene is expressed in the DA at low enough levels to compensate for the mutants’ deficits in *klf2* expression without inducing gross gain-of-function phenotypes. Indeed, this transgene’s expected equimolar co-expression of farnesylated mCherry is hard to visualize, consistent with limited expression levels. Another explanation is the existence of negative *klf2* autoregulation, in agreement with the semi-redundancy of the piscine *klf2* genes and data from cultured ECs showing that exogenous KLF2 represses the endogenous gene’s expression[30, 93].

At the cellular level, we find that Plxnd1, via its Klf2 mechano-effector, enlarges ECs as blood flow intensifies, thereby widening the DA during the 24-48 hpf period (see[66, 147]). The supporting evidence comes from morphometric analyses after this vessel’s expansion phase. First, *plxnd1^fov01b^* nulls and *klf2* mutants harbor supernumerary ECs with smaller areas and volumes within their reduced DAs. In contrast, *plxnd1^GAP1^* fish with appropriate DA caliber lack these cellular defects. Second, *sih* mutants without circulation feature a narrow DA with an endothelial surplus. While their junctional maturation defects hinder morphometric analyses based on the *TgKI(tjp1a-eGFP)^pd1252^* reporter, we deduce that their ECs are abnormally small. Third, forced endothelial expression of the WT receptor rectifies the narrow DA caliber in *plxnd1^fov01b^* nulls by increasing the area and volume of ECs and decreasing their abundance. However, forced endothelial expression of *klf2a* augments the DA caliber of both *plxnd1^fov01b^* nulls and *sih* mutants, enlarging the size of ECs without reducing cell numbers. The latter finding indicates that EC enlargement is sufficient for DA expansion and that ECs can change their size without altering their number. Fourth, consistent with the latter notion, in *gata1a^m651^* mutants with diminished circulatory shear stress, the circulation minimally increases the DA caliber during this vessel’s expansion phase. Yet, EC abundance during this period is similar between these mutants and their WT-like sibs.

Furthermore, our data contradicts alternative cellular mechanisms of DA caliber regulation. For instance, the model that the DA expands its caliber from 24-48 hpf through EC addition is inconsistent with our finding that in WTs and the two *plxnd1* mutants analyzed, the DA caliber expands as EC abundance diminishes, albeit this occurs at a slower pace in both mutants. The hypothesis that EC overabundance leads to DA narrowing in *sih*, *klf2*, and *plxnd1^fov01b^* mutants is irreconcilable with two observations. First, despite the *sih* mutant’s rising EC numbers, their narrow DA caliber remains constant. Second, forced endothelial expression of Klf2a in *sih* and *plxnd1^fov01b^* mutants enlarges the footprint and volume of ECs without affecting their abundance, expanding the DA caliber of these fish. Hence, although *sih*, *plxnd1*, and *klf2* mutants display elevated EC abundance, this cellular defect is not responsible for their reduced DA caliber.

However, it is unclear how lack of circulation, loss of Plxnd1 mechanosensing, and *klf2* inactivation increase the DA’s EC abundance. Perhaps the failure to increase EC size somehow enhances cellular proliferation, survival, or both[148, 149]. Alternatively, the flow-dependent specification and extrusion of Hematopoietic Stem Cells (HSCs) at the DA’s floor might cause these cells or their EC precursors to remain within the vessel or delay their exit[147, 150].

Our piscine and murine findings indicate that endothelial *PLXND1* is a positive regulator of the DA’s caliber across vertebrates. Additionally, using engineered MVNs made of human ECs carrying a transcriptional reporter of *KLF2* expression within a microfluidic chip, we found that flow induces statistically significant increases in the expression of the KLF2-GFP-reporter, vessel diameter, and EC size when compared to MVNs cultured under static (no flow) conditions. Remarkably, the differential effects of flow and static conditions in the MVNs parallel those observed in the piscine DA of WT embryos and *sih* mutants devoid of blood flow. The results from our experiments with vertebrate models and MVNs argue that circulatory forces exert similar molecular and cellular effects from fish to mammals. They also suggest that the receptor’s mechanosensing-dependent effects on vascular caliber regulation rely on evolutionarily conserved molecular and cellular mechanisms.

Although this study centers on the role of Plxnd1-mediated mechanosensing in regulating the DA caliber, our data suggests that the receptor might also determine the size of the trunk’s axial vein, the PCV. However, the relationship between Plxnd1 and flow is different in this case. Given its widespread endothelial expression[59, 68-70, 92, 151-157], *plxnd1* might exert a broad influence on the caliber of other vessels, including arteries, veins, capillaries with high shear stress levels (found in skeletal muscles[158] and the kidney’s glomeruli[159]), and lymphatic vessels[75, 76, 154]). In agreement with the notion that the effects of Plxnd1 on vascular caliber may be contextual, the cerebral vessels widen after transient brain infarction in mice with endothelial *Plxnd1* depletion[155].

Rescue experiments of *klf2* double mutant fish indicate that endothelial *klf2* activity also prevents the extrusion of ventricular cardiomyocytes in the heart[93], forecasting that Plxnd1-mediated mechanosensing may promote myocardial wall integrity by ensuring the abundance of this mechano-effector in ECs. Moreover, the central role of fluid shear stress in endocardial, venous, and lymphatic valve formation suggests that the Plxnd1-mediated detection of this mechanical stimuli regulates valvulogenesis across the cardiovascular system. Fittingly, both the receptor[68, 69, 71, 74, 92, 154, 160] and Klf2 express in the piscine and mammalian endocardium and orchestrate the morphogenesis of cardiac valve leaflets in fish and mice[161-164].

The bi-functional nature of the receptor prompts consideration of its activities’ regulation. While the flexibility of the Plxnd1 ectodomain seems necessary for mechanosensing[29], our findings imply that its intracellular GAP activity, key for guidance, is dispensable for sensing or interpreting circulatory forces. We envision that Plxnd1 carries out its distinct roles within separate sub-cellular compartments, with apical-junctional (see[29]) and basolateral (see[165, 166]) receptor pools mediating mechanosensing and guidance, respectively. This model implies that the proximal effectors and modulators of these distinct functions show a similar pattern of distribution or activation.

We anticipate that Plxnd1 harbors specific and shared molecular determinants of its two activities. Accordingly, mutations in this receptor might exert selective or pervasive functional effects. Human *PLXND1* variants are associated with congenital cardiac abnormalities, including defects in how blood vessels connect to the heart, often leading to the carriers’ fetal demise[167-170]. These abnormalities fit the phenotypes of mice with global or endothelial removal of *Plxnd1* and are attributable, at least partly, to the loss of its guidance function[28, 69- 71, 74, 154, 171]. However, some of these variants might additionally affect mechanosensing. Perhaps other *PLXND1* variants[172] selectively alter the ability of the receptor to detect blood flow forces. These considerations invite using model systems to uncover the molecular determinants of the receptor’s functions, verify the pathogenicity of *PLXND1* variants, and define the specificity of their functional impact.

The novel role of *PLXND1* and *KLF2* as vascular caliber regulators suggests that variants in these genes could contribute to diseases involving reduced vascular calibers, myocardial ischemia[173, 174], peripheral artery disease[175], atherosclerosis[29], hypertension[176], diabetes[177], and glaucoma[178]. Thus, increasing the expression or activity of Plexin D1 and KLF2 might therapeutically enhance the blood flow in individuals afflicted by these conditions and in patients with congenital heart defects suffering from impaired cardiac blood flow. This strategy could also potentiate some cancer treatments by improving the delivery of anti-tumor drugs and vascular-disrupting reagents[179, 180].

### Limitations of the study

**(1)** Our animal experiments employ receptor alleles that specifically lack guidance activity (this study) or are null[58, 59, 71], devoid of both guidance and mechanosensing functions. (**2**) Rescue experiments with 2D EC cultures argue that locking the receptor in its closed conformation (by introducing an intramolecular disulfide bond between the receptor’s first and ninth extracellular domains) abrogates its mechanosensing activity[29]. However, we have not tested the ability of such a form to rescue the DA caliber deficits of fish or mice lacking the endogenous receptor. (**3**) Besides eliminating cardiac contractility and RBCs, it is difficult to precisely manipulate piscine blood flow forces for an extended period[49, 50, 181, 182]. This challenge also applies to mice embryos. (**3**) The circulation deficiency of *sih* mutant fish is an extreme condition with potential secondary effects[183, 184]. However, as demonstrated by the *gata1a* mutants, reduced circulatory shear stress induces qualitatively similar effects on DA caliber. (**4**) The murine heart’s outflow tract arises as a single tube, the truncus arteriosus, that later septates into the aorta and the pulmonary artery. The septation fails in *Tie2^Cre/+^;Plxnd1^flox/flox^*embryos and mice are born with persistent truncus arteriosus. Some *Tie2^Cre/+^;Plxnd1^flox/flox^* embryos also develop ventricular septal defects. These congenital heart defects increase blood flow into the lungs, reducing irrigation elsewhere[71, 74]. However, it is unclear when these circulatory changes begin and whether they affect DA size. (**5**) We have not evaluated the impact of manipulating *PLXND1* and *KLF2* levels in MVNs under static or flow conditions. We highlight that the coherency of our zebrafish, mouse, and engineered MVN made of human ECs data fully supports our conclusions.

## SUPPLEMENTARY FIGURES

**Figure S1.**
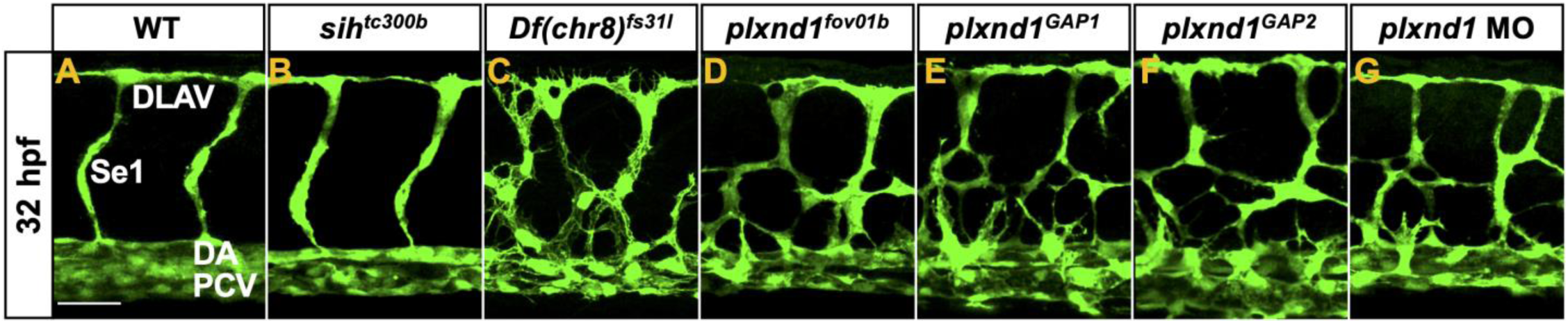
Angiogenic patterning in WTs, *sih* and *plxnd1* mutants, and *plxnd1* morphants at 32 hpf. Lateral views of the trunk’s vasculature in fixed, immune-fluorescently stained embryos of various genotypes. Endothelium (green), *Tg(fli1:EGFP)^y1^*. The dorsal side is up, and the anterior side is left. (**A**) WT. The scale bar (horizontal white line) is 40 μm units long. (**B**) *sih^tc300b^* mutant. (**C**) *plxnd1^Df(Chr08)fs31l^* mutant. (**D**) *plxnd1^fov01b^* mutant. (**E**) *plxnd1^GAP1^* mutant. (**F**) *plxnd1^GAP2^* mutant. (**G**) *plxnd1* morphant.

**Figure S2.**
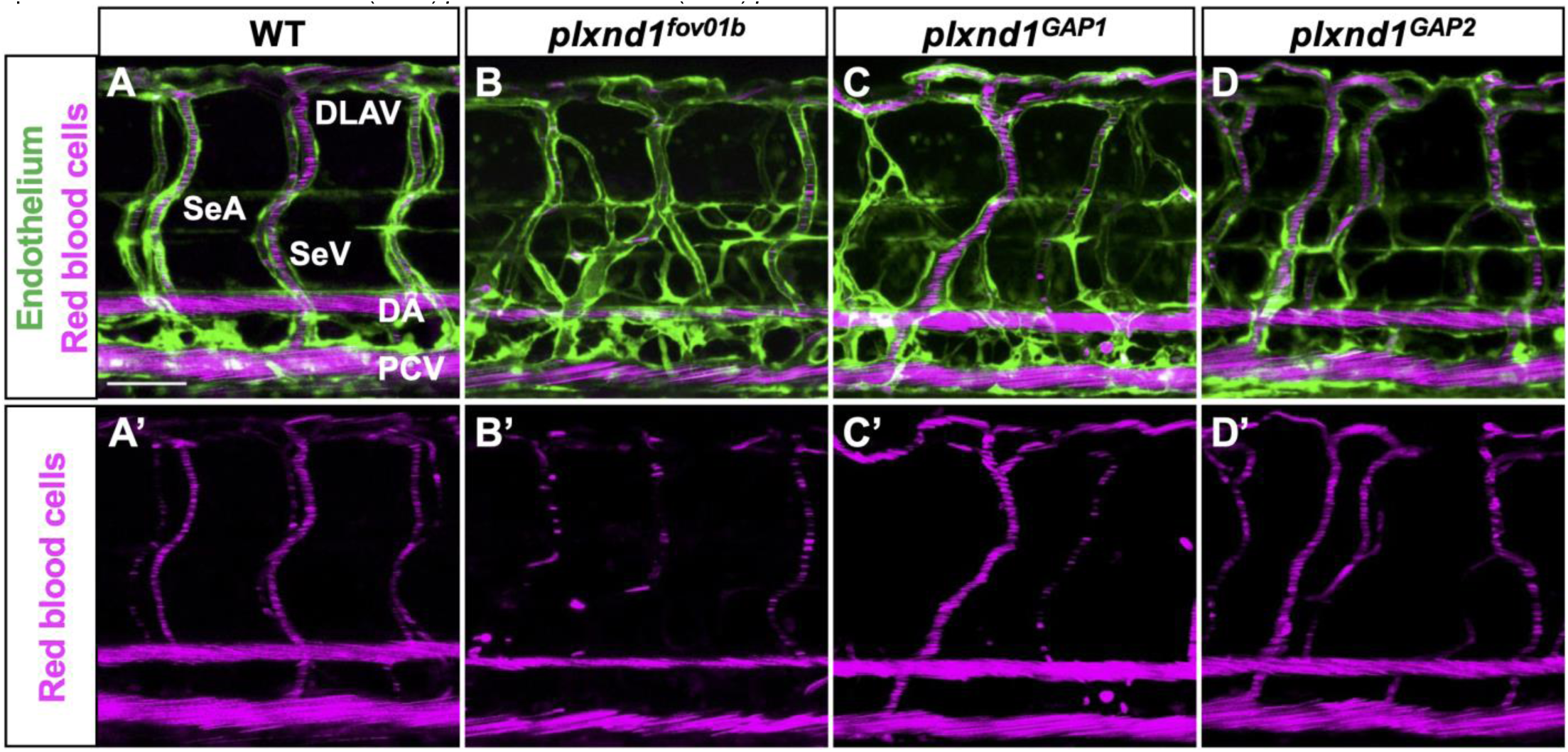
Red blood cell circulation in WTs and *plxnd1* mutants at 84 hpf. Lateral views of the trunk’s vasculature in live embryos. Endothelium (green), *Tg(fli1:EGFP)^y1^* reporter (A, B, C, D, E). Erythrocytes (magenta), *Tg(gata1a:dsRed)^sd2^* reporter[187] (A-E’). The fast-moving erythrocytes within the axial vessels appear as closely packed diagonal red lines. The dorsal side is up, and the anterior side is left. (**A-A’**) WT. The scale bar (horizontal white line) is 60 μm long. (**B-B’**) *plxnd1^fov01b^*. The erythrocytes occupy a narrower luminal space within the DA and PCV. (**C-C’**) *plxnd1^GAP1^* mutant. (**D-D’**) *plxnd1^GAP2^* mutant.

**Figure S3.**
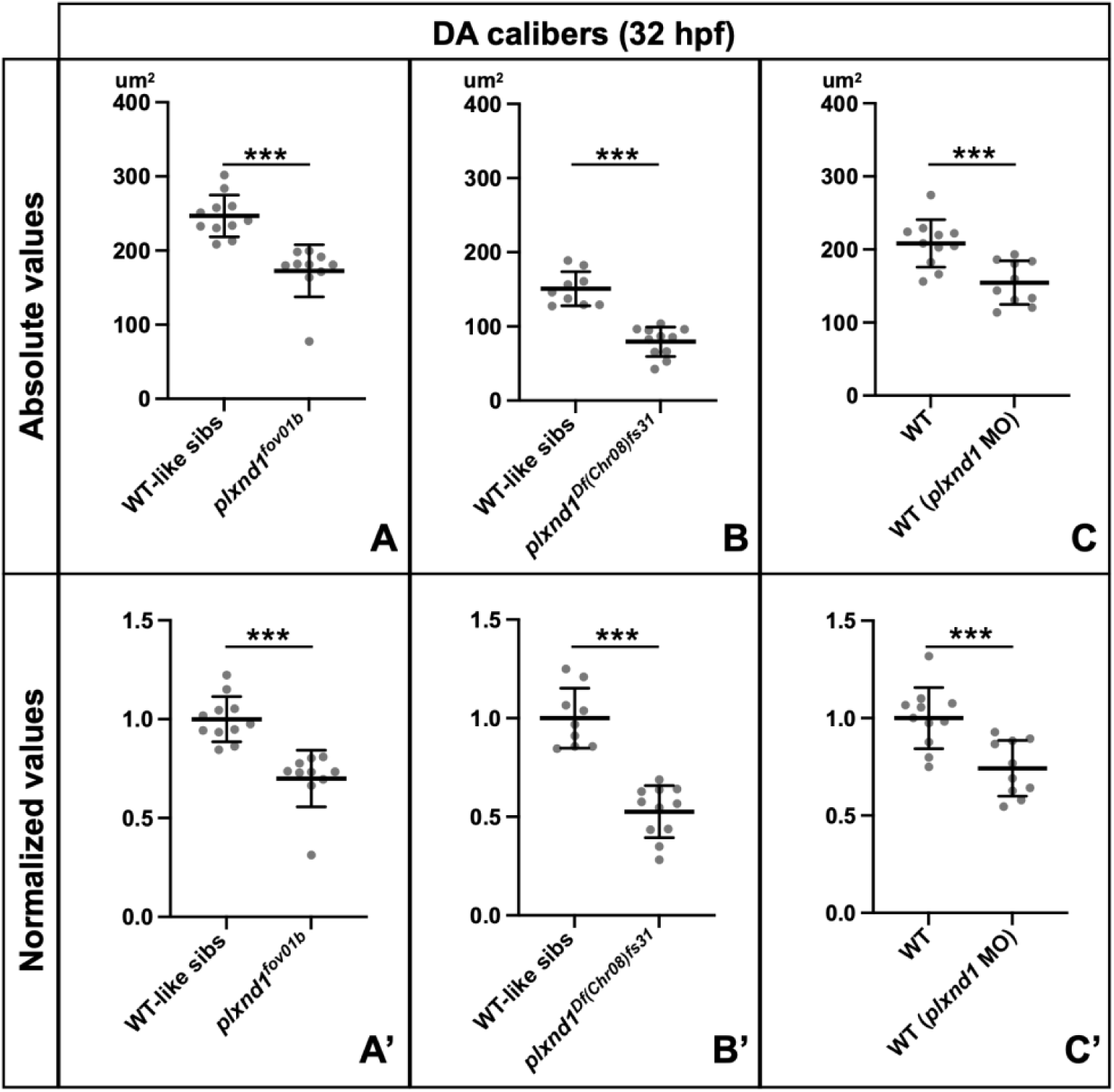
Both *plxnd1^fov01b^* and *plxnd1^Df(Chr08)fs31^* mutants and *plxnd1* morphants show reduced DA calibers at 32 hpf. (**A-C’**) DA (luminal area of DA cross-sections) in fixed siblings (sibs) of different genotypes. Statistical measures are as follows. The grey dots represent individual data points. The long and thick horizontal lines represent the means. The short and thin horizontal lines represent the error bars (SD). Significance levels as ∗p≤0.05, ∗∗p≤0.01, ∗∗∗p≤0.001, two-tailed unpaired Student’s *t*-test. (**A, B, C**) Comparison of absolute values. (**A’, B’, C’**) Comparison of normalized DA calibers. (**A-A’**) Comparison of WT-like sibs vs. *plxnd1^fov01b^* mutants. WT-like sibs group (n = 11): WTs and both single (*plxnd1^fov01b^/+* and *sih^tc300b^/+*) and double (*plxnd1^fov01b^/+*; *sih^tc300b^/+*) heterozygotes. *plxnd1^fov01b^* mutants (n = 10): *plxnd1^fov01b^* and *sih^tc300b^*/+; *plxnd1^fov01b^*. p <0.0001. (**B-B’**) Comparison of WT-like sibs vs. *plxnd1^Df(Chr08)fs31^* mutants. WT-like sibs group (n = 9): WTs and both single (*plxnd1^Df(Chr08)fs31^/+* and *sih^tc300b^/+*) and double (*plxnd1^Df(Chr08)fs31^/+*; *sih^tc300b^/+*) heterozygotes. *plxnd1^Df(Chr08)fs31^* mutants (n = 11): *plxnd1^Df(Chr08)fs31^* and *sih^tc300b^*/+; *plxnd1^Df(Chr08)fs31^*. p <0.0001. (**C-C’**) Comparison of WT vs. WT (*plxnd1* MO); p = X. WT group (n = 11): Un-injected WTs. WT (*plxnd1* MO) group (n = 10): WTs with *plxnd1* MO injection. p = 0.0009.

**Figure S4.**
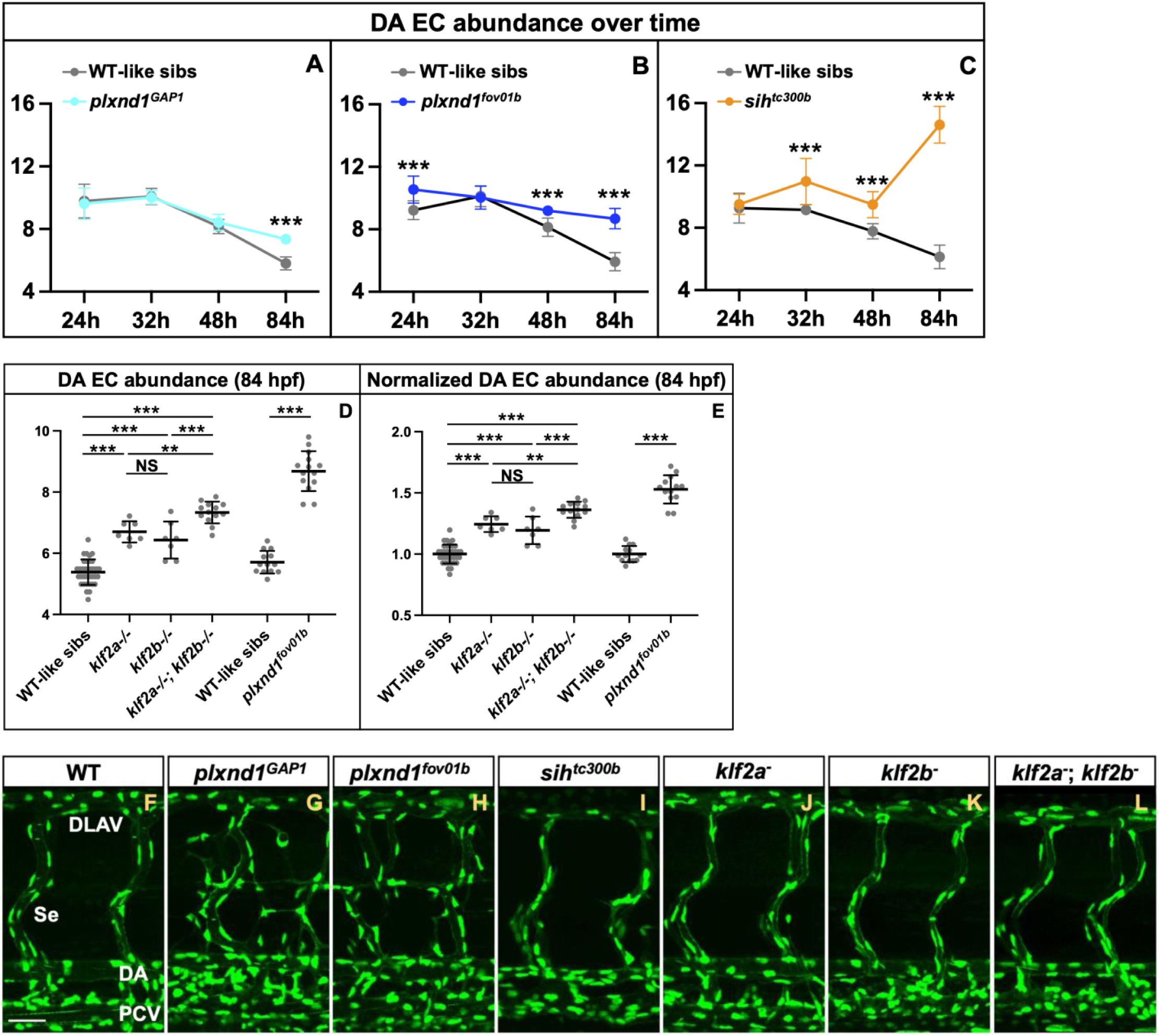
Comparison of EC abundance in the DA of live *plxnd1, sih*, and *klf2* mutants and their sibs. (**A-C**) Graphs comparing the DA’s EC abundance over time (at 24, 32, 48, and 84 hpf) in *plxnd1^GAP1^* (A), *plxnd1^fov01b^* (B), and *sih^tc300b^* mutants and their WT-like sibs (homozygous WTs and heterozygotes). Data is color-coded by genotype. Statistical measures are as follows: Dots denote each genotype’s means. Horizontal lines, error bars (SD). Significance levels as *p≤0.05, ∗∗p≤0.01, ∗∗∗p≤0.001); unpaired two-tailed Student’s *t*-test. (**A**) WT-like sibs (grey) vs. *plxnd1^GAP1^* mutants (cyan) comparisons are as follows. 24 hpf: WT-like sibs (n=12) vs. *plxnd1^fov01b^* (n=14), p=0.6883. 32 hpf: WT-like sibs (n=10) vs. *plxnd1^fov01b^* (n=9), p=0.8054. 48 hpf: WT-like sibs (n=9) vs. *plxnd1^fov01b^* (n=9), p=0.2542. 84 hpf: WT-like sibs (n=12) vs. *plxnd1^fov01b^* (n=12), p<0.0001. (**B**) WT-like sibs (grey) vs. *plxnd1^fov01b^* (navy blue) comparisons are as follows. 24 hpf: WT-like sibs (n=11) vs. *plxnd1^fov01b^*(n=15), p=0.0002. 32 hpf: WT-like sibs (n=13) vs. *plxnd1^fov01b^*(n=14), p=0.7443. 48 hpf: WT-like sibs (n=9) vs. *plxnd1^fov01b^* (n=9), p<0.0001. 84 hpf: WT-like sibs (n=13) vs. *plxnd1^fov01b^* (n=14), p<0.0001. (**C**) WT-like sibs (grey) vs. *sih^tc300b^* (orange) comparisons are as follows. 24 hpf: WT-like sibs (n=7) vs. *sih^tc300b^* (n=7), p=0.7043. 32 hpf: WT-like sibs (n=10) vs. *sih^tc300b^* (n=11), p=0.0012. 48 hpf: WT-like sibs (n=10) vs. *sih^tc300b^* (n=10), p=0.0001. 84 hpf: WT-like sibs (n=10) vs. *sih^tc300b^* (n=14), p<0.0001. (**D-E**) Graphs comparing the DA’s EC abundance (D) and the normalized (using the WT-like sibs as reference) DA’s EC abundance (E) at 84 hpf in both single (*klf2a^-^* and *klf2b^-^*) and double *klf2* (*klf2a^-^*; *klf2b^-^*) mutants, *plxnd1^fov01b^* mutants, and their WT-like sibs (homozygous WTs and single heterozygotes for both the single and double mutants). Statistical measures are as follows. Dots denote individual data points. Long horizontal lines represent the means. Short horizontal lines, error bars (SD). Significance levels as *p≤0.05, ∗∗p≤0.01, ∗∗∗p≤0.001), unpaired two-tailed Student’s *t*-test. WT-like sibs (n=35) vs. *klf2a^-^* (n=7), p<0.0001. WT-like sibs (n=35) vs. *klf2b^-^* (n=7), p<0.0001. WT-like sibs (n=35) vs. *klf2a^-^*; *klf2b^-^* (n=13), p<0.0001. *klf2a^-^* (n=7) vs. *klf2b^-^* (n=7), p=0.3244. *klf2a^-^* (n=7) vs. *klf2a^-^*; *klf2b^-^* (n=13), p=0.0012. *klf2b^-^* (n=7) vs. *klf2a^-^*; *klf2b^-^* (n=13), p=0.0005. WT-like sibs (n=13) vs. *plxnd1^fov01b^* (n=14), p<0.0001. (**F-L**) Representative lateral views of the trunk vasculature in live 84 hpf embryos of the indicated genotypes. Endothelial nuclei (green), *Tg(fli:nEGFP)^y7^* reporter[100]. The dorsal side is up, and the anterior side is left. The scale bar (horizontal white line) represents 40 μm (F).

**Figure S5.**
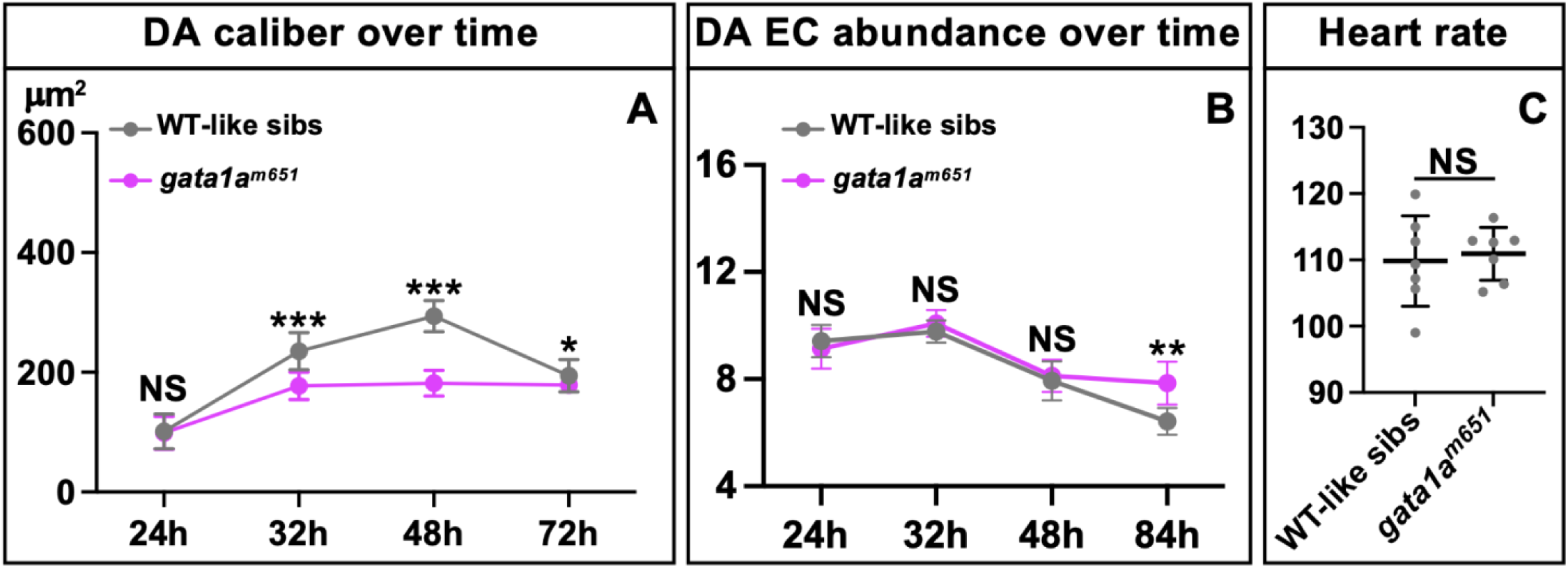
RBC-deficient *gata1a^m651^* mutants show reduced DA caliber but normal EC abundance and heartbeat frequency during this vessel’s circulation-induced expansion phase. (**A-C**) Comparisons of WT-like sibs (WTs and *gata1a^m651^*/+ heterozygotes) vs. *gata1a^m651^* mutants. Significance levels as NS>0.05, *p≤0.05, ∗∗p≤0.01, ∗∗∗p≤0.001; unpaired two-tailed Student’s *t*-test. (**A**) Aortic caliber in fixed embryos (luminal area of aortic cross-sections) at different time points (24, 32, 48, and 72 hpf). The dots represent means per genotype, and the short horizontal lines denote error bars (SD). WT-like sibs (grey) vs. *gata1a^m651^* mutants (magenta). WT-like sibs per stage: 24 hpf (n = 13), 32 hpf (n = 10), 48 hpf (n = 11), 72 hpf (n = 12). *gata1a^m651^* mutants per stage: 24 hpf (n = 10), 32 hpf (n = 12), 48 hpf (n = 9), 72 hpf (n = 14). Significance levels per stage: 24 hpf (p=0.8284), 32 hpf (p<0.0001), 48 hpf (p<0.0001), 72 hpf (p=0.0126). (**B**) aEC abundance at different time points (24, 32, 48, and 84 hpf) in live embryos. The dots represent means per genotype, and the short horizontal lines denote error bars (SD). WT-like sibs (grey) vs. *gata1a^m651^* mutants (magenta). WT-like sibs per stage: 24 hpf (n = 12), 32 hpf (n = 7), 48 hpf (n = 12), 72 hpf (n = 7). *gata1a^m651^* mutants per stage: 24 hpf (n = 5), 32 hpf (n = 7), 48 hpf (n = 8), 72 hpf (n = 6). Significance levels per stage: 24 hpf (p=0.4110), 32 hpf (p<0.0001), 48 hpf (p<0.0001), 72 hpf (p=0.0126). (**C**) Heart rate at 50 hpf. Gray dots represent data from individual embryos, the wide horizontal lines are means, and the short horizontal lines represent error bars (SD). WT-like sibs: n=7. *gata1a^m651^*: n=7. Significance level: p=0.7220.

**Figure S6.**
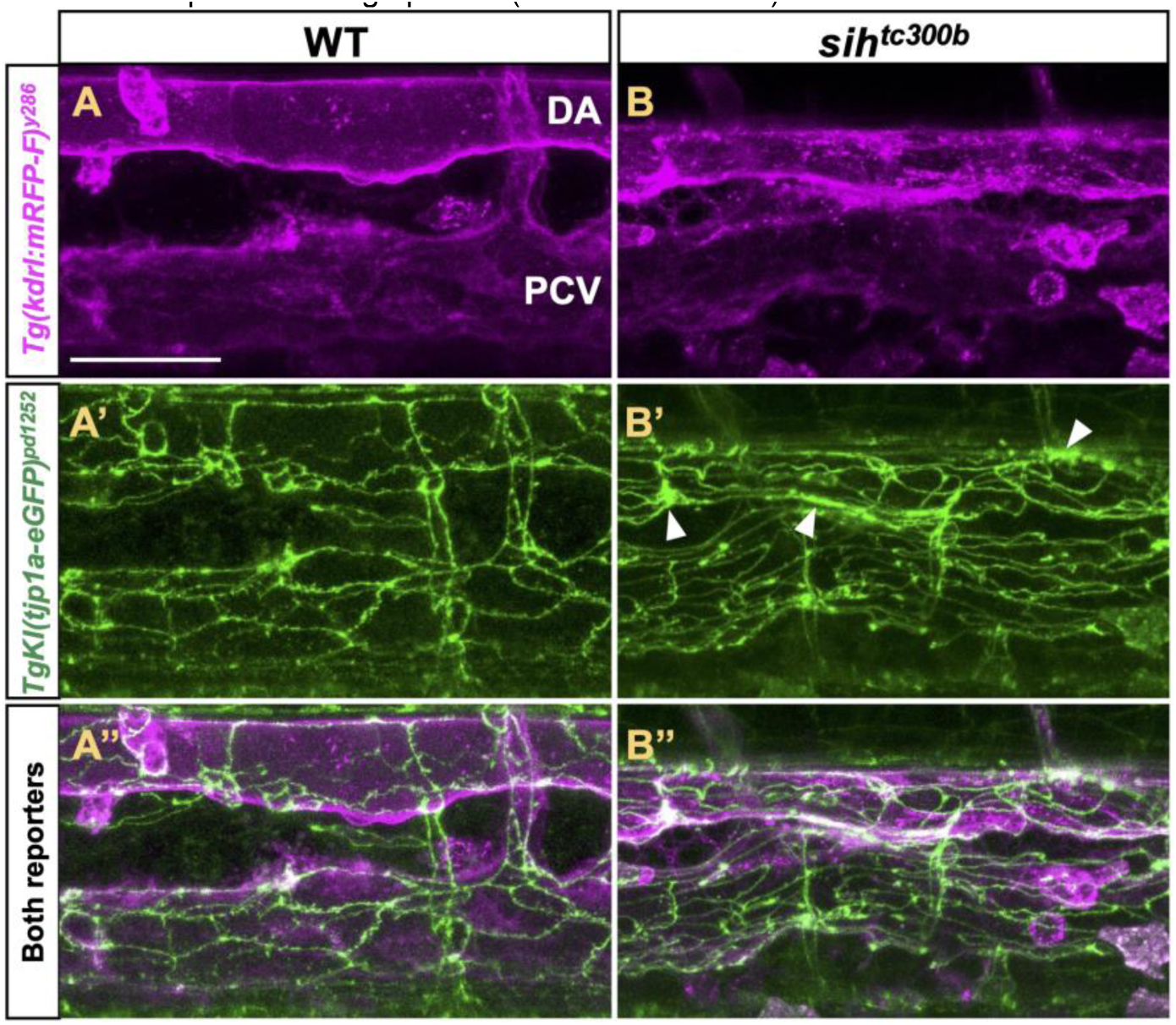
Endothelial junction defects in *sih* mutants. The panels show lateral views of the trunk’s axial vessels from live 84 hpf embryos. The dorsal side is up, and the anterior side is left. (**A, B**) Endothelial membranes (red), *Tg(kdrl:mRFP-F)^y286^*. (**A’, B’**) Endothelial Tjp1a-eGFP (+) tight junctions (green), *TgKI(tjp1a-eGFP)^pd1252^*. (**A’’, B’’**) Both reporters. (**A-A’’**) WT-like sib. The scale bar (horizontal white line) is 40 μm. (**B-B’’**) *sih^tc300b^* mutant. Note that the DA shows tight junctions of variable widths and accumulation of the Tjp1a- eGFP fusion protein in large puncta (white arrowheads).

**Figure S7.**
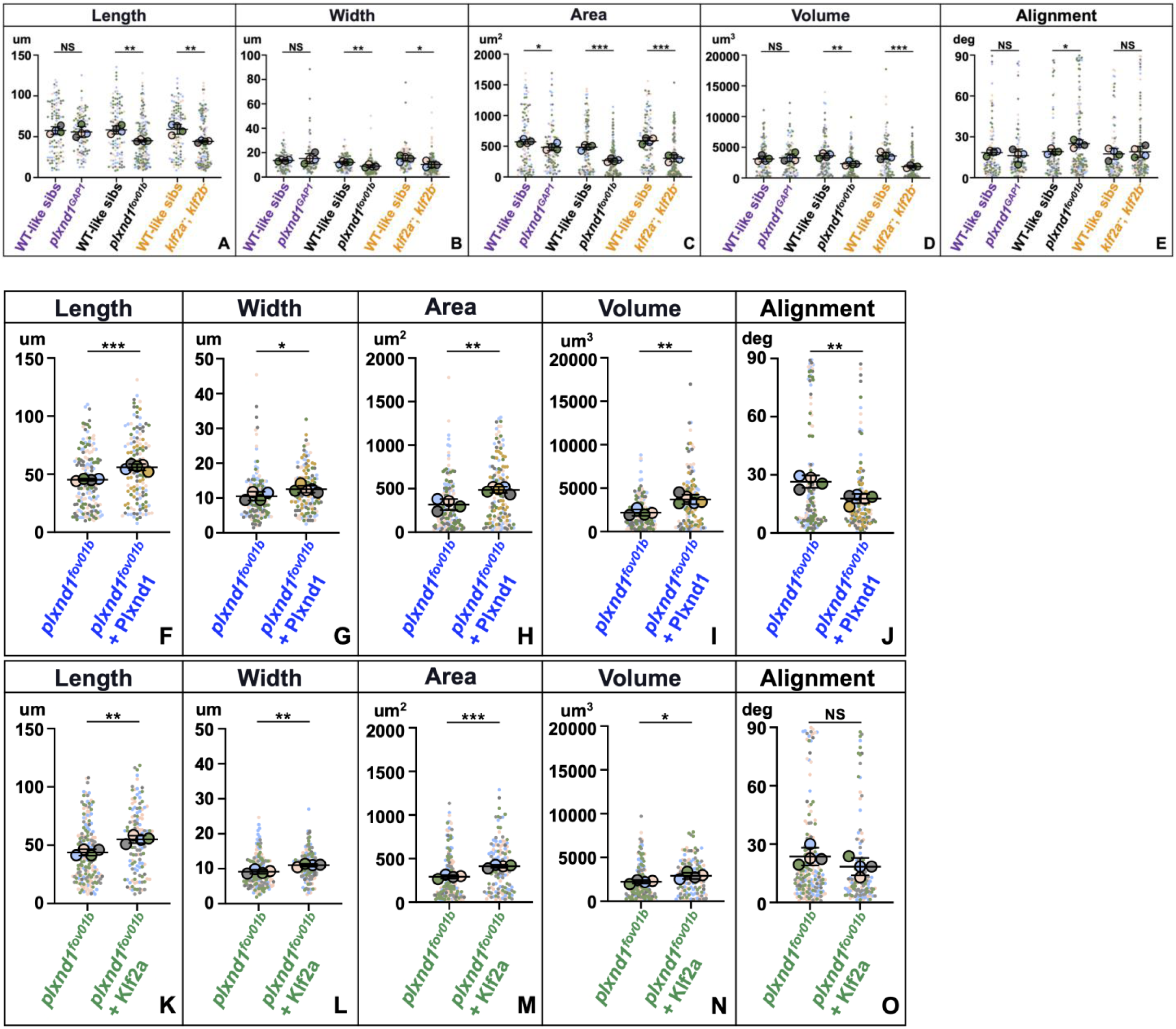
Plxnd1 expands the DA caliber by enlarging the area of ECs via Klf2. (**A-O**) Superplot graphs[186] compare the EC morphometry of live mutants’ and their WT-like sibs: length (A, F, K), width (B, G, L), area (C, H, M), volume (D, I, N), and alignment angle (E, J, O). The mean values for all these genotypes are also in Figure 4, except for *plxnd1GAP1* and their WT-like sibs, shown here only (see below). <u>Statistical measures are as follows</u>. Large circles denote means color-coded for each larva. Small dots represent data from individual ECs, color-coded for each larvae. The long, thick horizontal lines represent the means. Short and thin horizontal lines, error bars (SD). Significance levels as *p≤0.05, ∗∗p≤0.01, ∗∗∗p≤0.001), unpaired two-tailed Student’s *t*-test. (**A-E**) Comparisons of *plxnd1^GAP1^*, *plxnd1^fov01b^*, and *klf2* (*klf2a^-^*; *klf2b^-^*) double mutants against their WT-like sibs (homozygous WTs for the three mutants and heterozygotes for the *plxnd1^GAP1^* and *plxnd1^fov01^* alleles). WT-like sibs (4 embryos, 104 cells) vs. *plxnd1^GAP1^* (4 embryos, 105 cells), purple label: (A) p=0.6718, (B) p=0.4635, (C) p=0.0355, (D) p=0.6753, (E) p=0.4348. WT-like sibs (4 embryos, 101 cells) vs. *plxnd1^fov01b^* (4 embryos, 162 cells), black label: (A) p=0.0018, (B) p=0.0094, (C) p=0.0001, (D) p=0.0022, (E) p=0.0128. WT-like sibs (4 embryos, 103 cells) vs. *klf2* double mutants (4 embryos, 148 cells), orange label: (A) p=0.0031, (B) p=0.0231, (C) p=0.0003, (D) p=0.0009, (E) p=0.6077. (**F-J**) Comparisons of *plxnd1^fov01b^* (4 embryos, 170 cells) vs. *plxnd1^fov01b^* + Plxnd1 (*Tg(fliep: 2xHA-plxnd1)*) (5 embryos, 133), blue label: (F) p=0.0003, (G) p=0.0041, (H) p=0.0003, (I) p=0.0026, (J) p=0.0039. (**K-O**) Comparisons of *plxnd1^fov01b^* (4 embryos, 129 cells) and *plxnd1^fov01b^* + Klf2a (*Tg(fliep:klf2a)*) (4 embryos, 97 cells), green label: (K) p=0.0018, (L) p=0.0016, (M) p=0.0001, (N) p=0.0180, (O) p=0.1613.

**Table S1.**
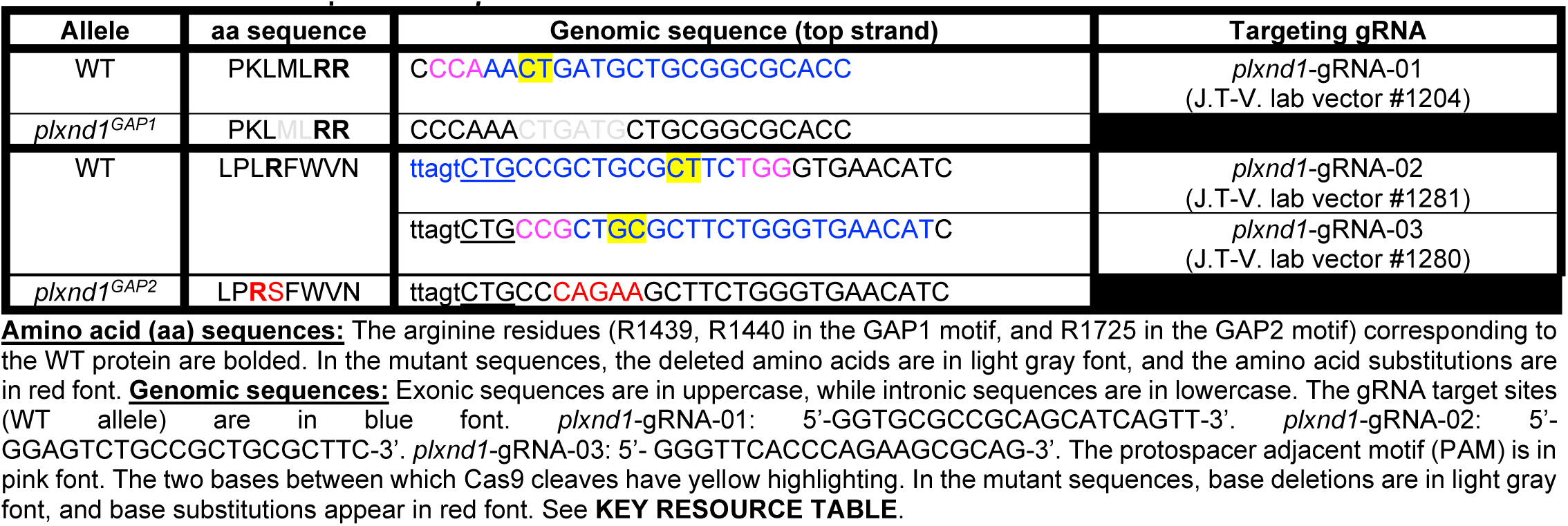
Molecular description of the *plxnd1^GAP^* mutant alleles.

**Table S2.**
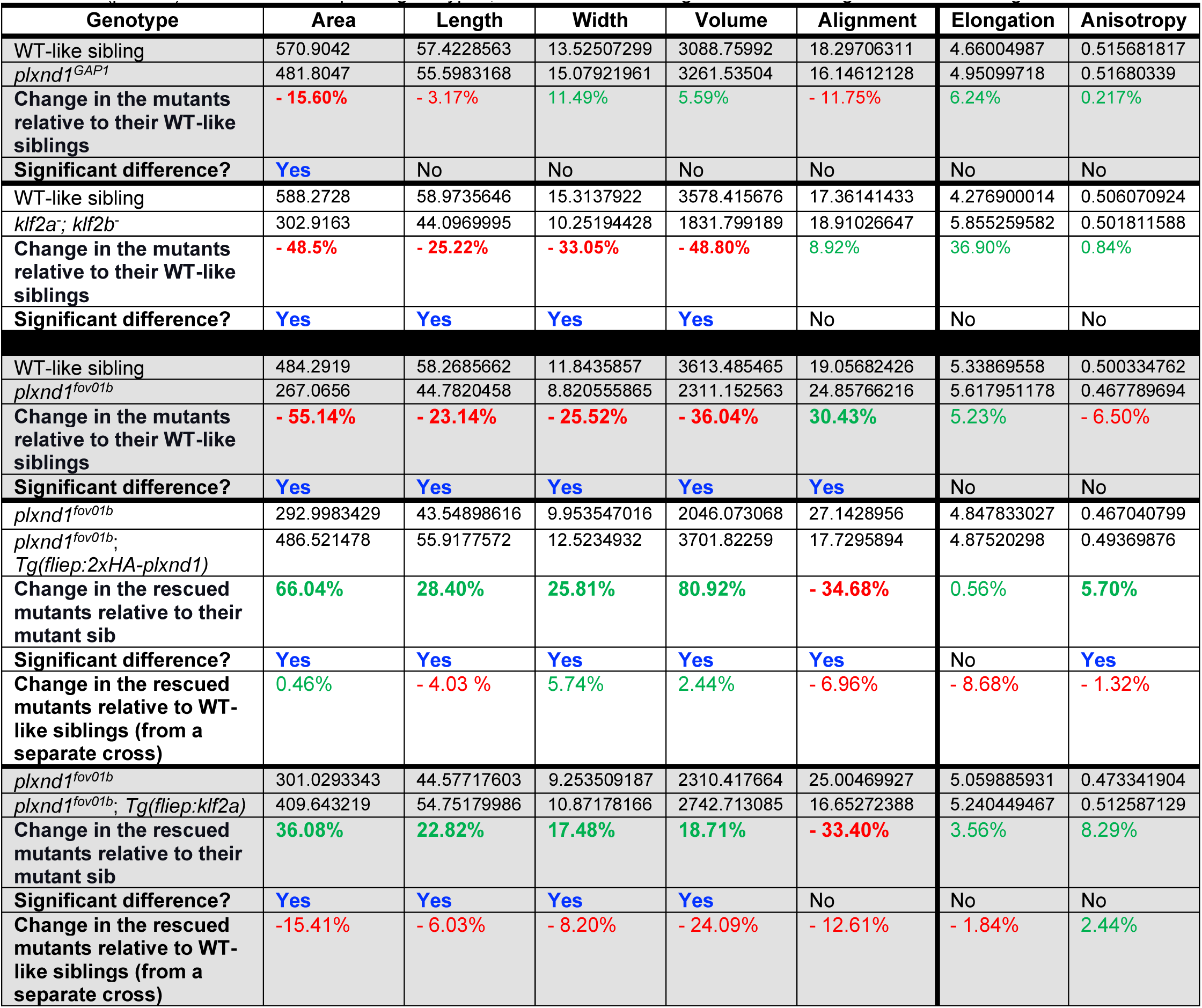
Comparisons of EC morphometry values (means of means) between genotypes at 84 hpf. The EC area, length, width, volume, and alignment values are the same as those in **Figures 4** and **S7**. The blue text denotes significant differences (p<0.05) between the compared genotypes, with red text denoting a reduction and green text indicating an increase.

## Funding

The NIH’s NHLBI grant 1R01HL161090-01A1 (to JTV) supported the murine, MVNs, and zebrafish studies in the MKS, GGC, and JTV labs. Grants from the Singapore Ministry of Health’s NMRC (MOH- OFIRG18nov-0036) and the Singapore Ministry of Education (MOE-T2EP30121-0025) also funded the work in the MKS lab. The NIH grants R21HL152367 and T32EB016652 (to GG-C and AB) sponsored the experiments with MVNs, too. The NIH grant 1R01DK1321120 (to MB) sponsored the generation of *TgKI(tjp1a-eGFP)^pd1252^*).

## Reagents

<u>Fixed embryos</u>: Sarah Childs (*plxnd1^Df(Chr08)fs31l^*;*Tg(kdrl:GFP)^la116^*and their WT-sibs)[97]. <u>Fish lines</u>: Brant Weinstein and Dan Castranova (*Tg(kdrl:mRFP-F)^y286^*)[117], Neil Chi and Julien Vermot (*Tg(klf2a:H2b- EGFP)^ig11^*)[87], Amber N. Stratman (*klf2a^y616^*)[86]. Michel Bagnat (*TgKI(tjp1a-eGFP)^pd1252^*)[112], the Zebrafish International Resource Center/ZIRC (*gata1a^m651^*, *klf2b^sa43252^*, and *sih^tc300b^*). <u>Vectors</u>: Chi-Bin Chien and Koichi Kawakami (pCS2FA-transposase, pDestTol2CG2, and p3E-polyA,)[188], Nathan Lawson (p5E-fliep)[189], Salim Abdelilah-Seyfried (pME-klf2a)[94], and Xingxu Huang (pST1374-NLS-flag-linker-Cas9; Addgene plasmid #44758)[190].

## Microscopy

Zebrafish confocal imaging was done at the NYU Grossman School of Medicine’s Microscopy Laboratory (partially supported by the NIH’s NCI grant P30CA016087) using a Leica SP8 confocal system (supported by the NIH’s NCRR grant 1S10RR024708). We thank Michael Cammer for imaging guidance.

## Scientific advice

We thank our colleagues E. Jane Albert Hubbard, Holger Knaut (and members of his lab), Ruth Lehman, Jeremy F. Nance, Niels Ringstad, Agnel Sfeir, Hyung Don Ryoo, and Jessica E. Treisman for useful suggestions.

## AUTHOR CONTRIBUTIONS

**AB:** Investigation, data curation, formal analysis, methodology, visualization, writing – review and editing. **DSL:** Resources, writing – review and editing. **GGC:** Supervision, writing-original draft, writing – review, and editing. **JH:** Resources, data curation, formal analysis, investigation, methodology, visualization, writing-original draft, Writing – review and editing. **JGR:** Software, writing-original draft, writing – review, and editing. **JTV:** Conceptualization, funding acquisition, project administration, supervision, resources, methodology, writing-original draft, writing – review and editing. **JM:** Resources, writing – review and editing. **MB:** Resources, writing – review and editing. **MKS:** Formal analysis, writing-original draft, writing – review and editing. **SCP:** Resources, writing – review and editing. **UT**: Data curation, formal analysis, methodology, visualization.

## METHODS

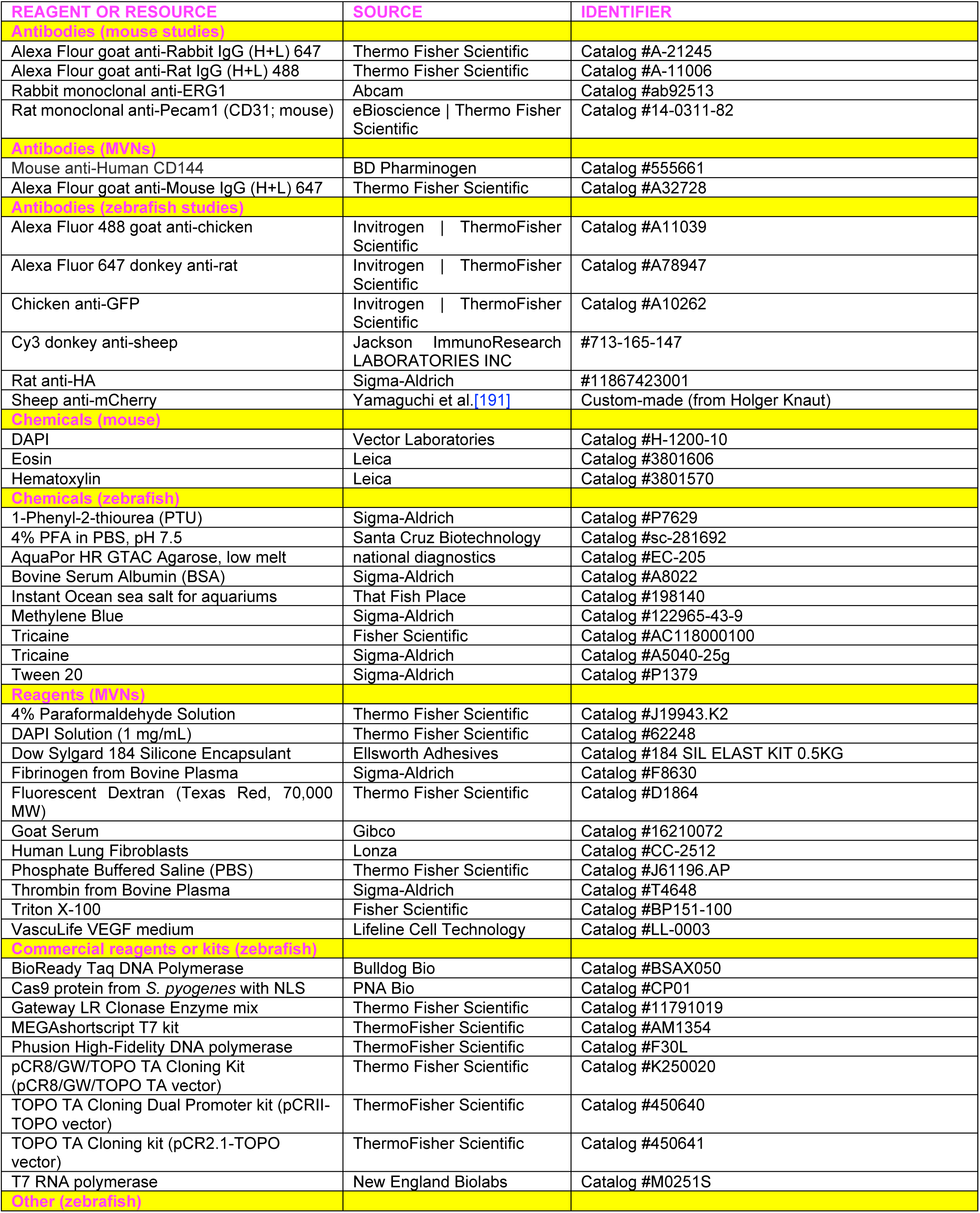
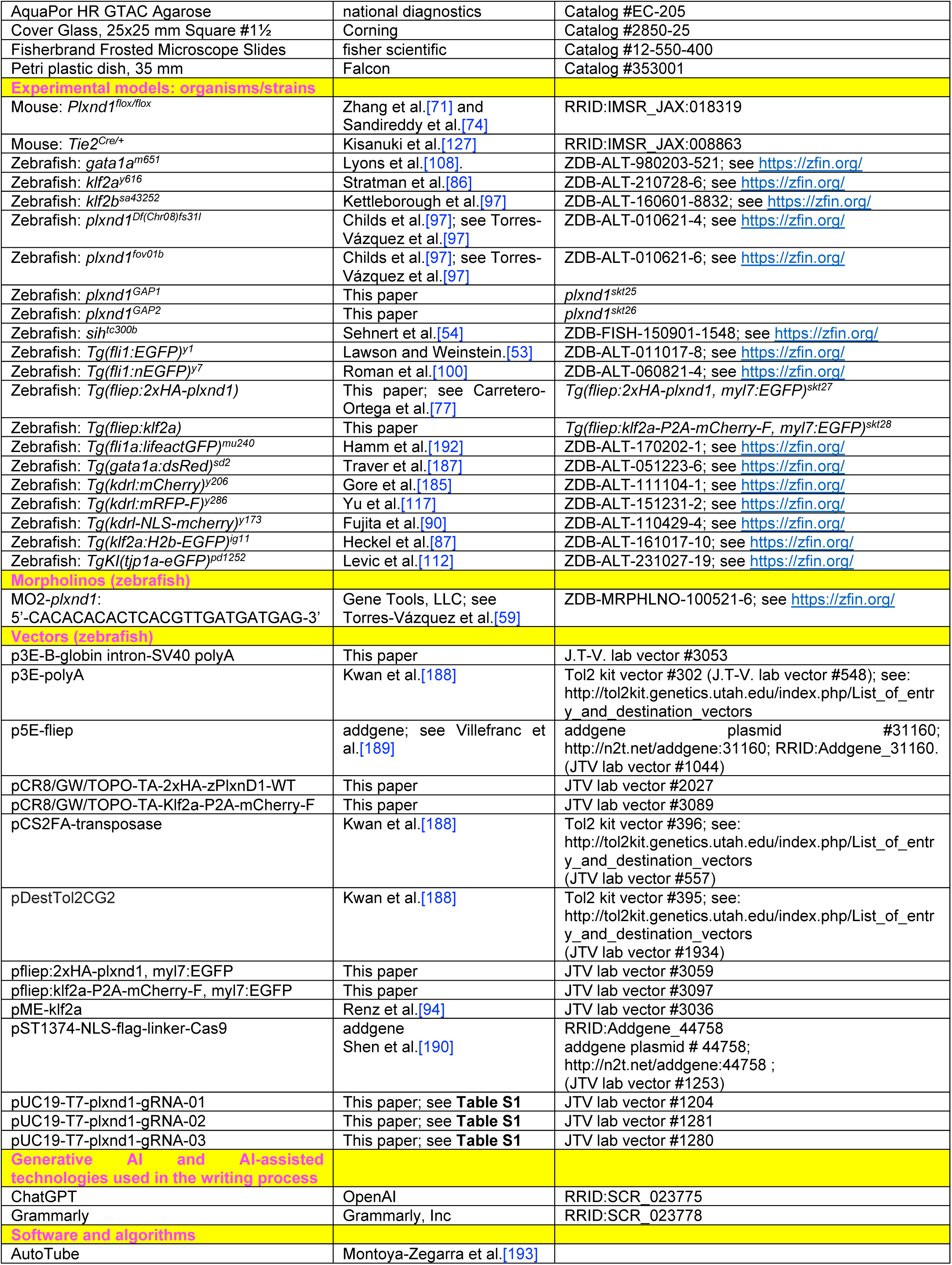
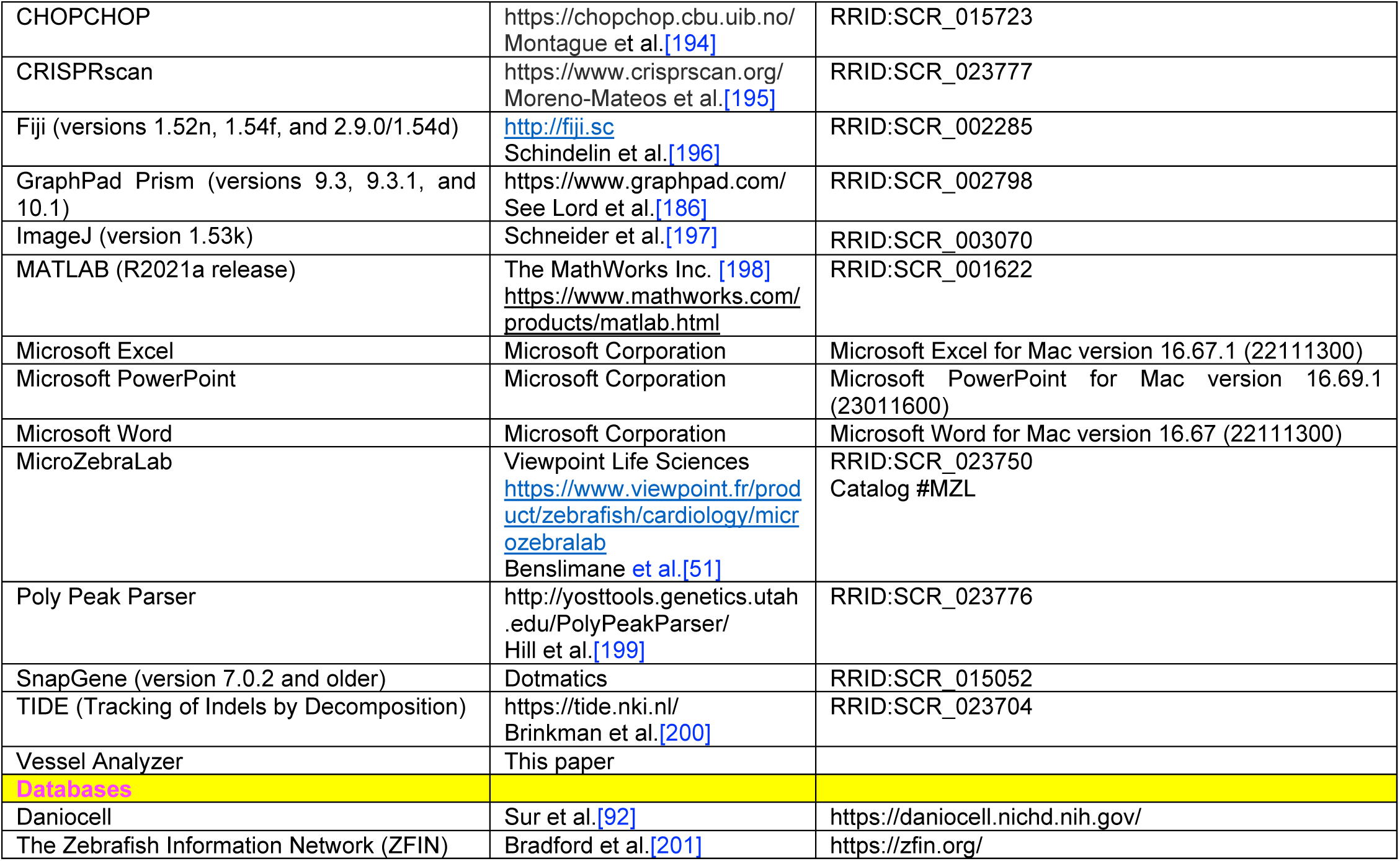
KEY RESOURCE TABLE.

## EXPERIMENTAL MODELS AND SUBJECT DETAILS

### Animals

#### Fish

We raised zebrafish (*Danio rerio*) in recirculating water systems (28°C, 14-hour light/10-hour dark cycle) at the NYU Grossman School of Medicine, following IACUC-approved animal protocols and under Veterinarian supervision (AAALAC International accredited). We bred fish in mating tanks, raising embryos to 5 dpf in Blue Water (comprising 20 L distilled water, 20 ml Methylene Blue aqueous solution (1g/L), and 6 grams of Instant Ocean sea salt for aquariums) or zebrafish system water (ZSW, the water in the aquaria) in a 28.5°C incubator. To prevent embryo pigmentation, we administered 1-Phenyl-2-thiourea (PTU) treatment before 24 hpf[202]. No tests were conducted on the influence of sex since sex determination occurs at later stages. <u>Mutant alleles</u>: *gata1a^m651^*(putative null) contains a C-to-T transversion at position 1015 of the open reading frame, yielding a predicted nonsense mutation at Arg339, truncating the last 79 aa of the Gata1a protein, including part of the basic domain critical for DNA binding[108]. *klf2a^y616^*(putative null) contains an eight bp insertion in exon 2, resulting in a frameshift and early termination, encoding an 181 aa protein with the N-terminal 157 residues of the 380 aa WT product[86] (transactivation domain and trans-repression domain fragment; see[93]). *klf2b^sa43252^*(putative null)[97] harbors a C-to-A transversion at position 32 of exon 1, converting Ser11 into a stop codon and truncating 353 aa. *plxnd1^Df(Chr08)fs31l^*(null) is a chromosomal deficiency of *plxnd1* and nearby genes[58, 59]. *plxnd1^fov01b^* (null) harbors a C-to-A transversion converting Tyr318 to a stop codon, truncating the 1,880 aa receptor early in its N-terminal extracellular Sema domain. Reduced mutant message expression is likely due to nonsense mRNA decay[58-60]. *plxnd1^GAP1^* (*plxnd1^skt25^*) and *plxnd1^GAP2^* (*plxnd1^skt26^*) delete and replace two amino acids in the receptor’s GAP1 and GAP2 motifs[26, 62-64] (this study, **Table S1**), likely eliminating the receptor’s GAP function. *sih^tc300b^*(null) features an A-to-G transition at the −2 position of the splice-acceptor in the second intron of *tnnt2a*, causing a frameshift and a premature stop codon in exon 7. The mutant message encodes only the first 11 aa of the WT protein[54]. <u>Transgenic lines</u>: *TgBAC(tjp1a-eGFP)^pd1252^*[112], *Tg(fli:nEGFP)^y7^*[100], *Tg(fli1:EGFP)^y1^*[53], *Tg(fli1a:lifeactGFP)^mu240^*[192], *Tg(fliep:2xHA-plxnd1)* and *Tg(fliep:klf2a)* (this study), *Tg(kdrl:mCherry)^y206^*[185], *Tg(kdrl:mRFP-F)^y286^*[117], *Tg(kdrl:nls-mCherry)^y173^*[90], and *Tg(klf2a:H2b- EGFP)^ig11^*[87]. Consult the **KEY RESOURCE TABLE**.

#### Mice

We used the *Tie2^Cre^* endothelial-specific Cre line[127] and the floxable *Plxnd1* allele *Plxnd1^flox^*[71, 74]. See the **KEY RESOURCE TABLE**.

### Engineered microvascular networks (MVNs) made from human ECs and application of flow

We fabricated microfluidic chips[203, 204] and pumps[205] from Polydimethylsiloxane (PDMS, Dow Corning Sylgard 184, Ellsworth Adhesives), assembling them as described before. We formed MVNs by suspending 7 million/ml HUVECs (Human Umbilical Vein Endothelial Cells; isolated and cultured as in[206] and carrying a flow-inducible reporter of *KLF2* transcription[123] and 1 million/ml of human lung fibroblasts (Lonza) in 1.4 U/mL of thrombin (Sigma-Aldrich) and 3 mg/mL of fibrinogen (Sigma-Aldrich) and injecting the cell suspension into the microfluidic chips. We maintained the resulting devices in VascuLife VEGF medium but with only a quarter (0.19 U/mL) of the heparin sulfate provided in the kit (Lifeline Cell Technology), changing the medium daily. We applied constant flow, resulting in a mean shear stress of 0.5 Pa in the MVNs using the microfluidic pump starting on day 5 of the culture. We maintained the flow for 48 hours before evaluating the MVNs. We did not expose MVNs cultured under static conditions to flow. We provide further details about these protocols in[124]. See also [125, 126]. Consult the **KEY RESOURCE TABLE**.

## METHOD DETAILS

### Animal genotyping

We amplified PCR products from DNA isolated from zebrafish (adult fin clips or whole embryo s or larvae) and murine (embryo yolk sacs and adult tail biopsies) lysates, using BioReady Taq DNA polymerase for the former. We analyzed the PCR products by agarose gel electrophoresis, Sanger sequencing, or both. We handled and annotated zebrafish allele sequences using SnapGene software.

### Zebrafish: genome editing

We generated *plxnd1^GAP^* alleles using CRISPR/Cas9-based genome editing[61], as in[77]. We selected gRNA targets using CHOPCHOP[194] and CRISPRscan[195] web tools. We used T7 RNA polymerase or the MEGAshortscript T7 kit to *in vitro* transcribe gRNAs from the pUC19-T7-plxnd1-gRNA-01, pUC19-T7-plxnd1-gRNA-02, and pUC19-T7-plxnd1-gRNA-03 vectors (made via PCR-assembly of oligo templates (Integrated DNA Technologies)[62, 207] subcloned into pUC19-T7[207]). We induced editing by delivering 1 nl of a mix of gRNA

(125 pg) and *S. pyogenes* Cas9 nuclease mRNA (300 pg; transcribed from pST1374-NLS-flag-linker-Cas9[190]) or protein (500 pg; PNA Bio) into one-cell stage *Tg(fli1a:EGFP)^y1^*[53] G0 embryos. We pooled genomic DNA from 4–12 G0s to assess genome editing, generating PCR amplicons with BioReady Taq DNA polymerase. We analyzed PCR products by agarose gel electrophoresis or sequencing (after TOPO-TA cloning into pCRII-TOPO or pCR2.1-TOPO vectors, with 8-20 colonies evaluated). We identified adult G0s with germline mutations via individual outcrossing with *plxnd1^fov01b^*; *Tg(fli1a:EGFP)^y1^* mutants, scoring the progeny for vascular misguidance. We outcrossed carrier G0s to WT *Tg(fli1a:EGFP)^y1^* fish to create F1 fish. We identified F1 heterozygous carriers via PCR genotyping and analysis of the sequence trace using the Poly Peak Parser[199] and TIDE[200] web tools. We confirmed allele sequences by single-colony Sanger sequencing of TOPO-TA cloned PCR products amplified from F1 genomic DNA. See **Table S1** and the **KEY RESOURCE TABLE**.

### Zebrafish: constructs for making transgenic lines

To make the *Tg(fliep:2xHA-plxnd1)* (full name: *Tg(fliep:2xHA-plxnd1, myl7:EGFP)^skt27^*) and *Tg(fliep:klf2a)* (full name: *Tg(fliep:klf2a-P2A-mCherry-F, myl7:EGFP)^skt28^* lines, we assembled the J.T-V. lab vectors #3059 (pfliep:2xHA-plxnd1, myl7:EGFP)[77] and #3097 (pfliep:klf2a-P2A-mCherry-F, myl7:EGFP) via Gateway cloning[208] using the Gateway LR Clonase Enzyme mix. We recombined the pDestTol2CG2 destination vector (with myl7:EGFP cardiac marker)[188], the entry clone p5E-fliEP (for endothelial expression)[189], and these other entry clones. For #3059: J.T-V. lab vectors #2027 (pCR8/GW/TOPO-TA-2xHA-zPlxnD1-WT; a pCR8/GW/TOPO TA derivative) and #3053 (p3E-B-globin intron-SV40 polyA). The latter includes the rabbit β- globin intron and the SV40 late polyA signal to promote mRNA export and enhance protein expression[209-213], enabling efficient expression of ∼6 kb cDNAs. For #3097, J.T-V. lab vectors #3089 (pCR8/GW/TOPO-TA-Klf2a- P2A-mCherry-F-P2A-mCherry-F and p3E-polyA[188]. Vector #3089 is a derivative of pCR8/GW/TOPO TA and pME-klf2a[94] made for P2A-based equimolar co-expression[214] of piscine Klf2a and farnesylated mCherry. J.T-V. lab vectors #2027 and #3089 harbor the optimal 5’ zebrafish KOZAK sequence (5’-GCAAAC-3’)[215] upstream of the 2xHA-zPlxnD1-WT (5,709 bp) and Klf2a (1,140 bp) ORFs to enhance expression. We designed and annotated plasmids using SnapGene software. We amplified PCR products for plasmid construction with Phusion High-Fidelity or BioReady Taq DNA polymerases. We made our entry clones via Gibson assembly[216]. See the **KEY RESOURCE TABLE.**

### Zebrafish: transgenesis

We performed *Tol2*-based transgenesis[217] with the J.T-V. lab vectors #3059 (pfliep:2xHA-plxnd1, myl7:EGFP) and #3097 (pfliep:klf2a-P2A-mCherry-F, myl7:EGFP) to make the *Tg(fliep:2xHA-plxnd1)* (full name: *Tg(fliep:2xHA-plxnd1, myl7:EGFP)^skt27^*) and *Tg(fliep:klf2a)* (full name: *Tg(fliep:klf2a-P2A-mCherry-F, myl7:EGFP)^skt28^* lines. We injected one nl of a solution containing 100 pg of Tol2 mRNA (synthesized *in vitro* from the pCS2FA-transposase vector[188]) and 200 pg of the transgenesis vector into the cytoplasm of one-cell stage *plxnd1^fov01b^* mutants carrying the *Tg(kdrl:mCherry)^y206^*, *Tg(fli:nEGFP)^y7^*, or *Tg(fli1a:lifeactGFP)^mu240^* endothelial reporters[100, 185, 192]. We scored G0 embryos at 1-3 dpf, raising the animals with cardiac green fluorescence. To make F1 lines, we outcrossed the G0s with *plxnd1^fov01b^* mutants, selecting fish with green hearts. We established F2 lines from single F1 fish. Expression of the farnesylated mCherry from the *Tg(fliep:klf2a)* line is usually undetectable by confocal microscopy in 1-3 dpf fish. See the **KEY RESOURCE TABLE**.

### Zebrafish: immunofluorescence

Embryos were fixed overnight (ON) at 4°C in 4% paraformaldehyde (PFA) solution in phosphate-buffered saline (PBS), pH 7.5. The embryos were then washed six times (10 min/wash) with PBST (PBS with 0.2% Tween) and incubated in the blocking solution (PBST with 1% bovine serum albumin (BSA)) for 1 hr, all at room temperature (RT). Embryos were then incubated ON at 4°C with primary antibodies diluted in blocking solution and then washed with PBST six times (10 min/wash) at RT. The embryos were then incubated overnight at 4°C with fluorescent secondary antibodies diluted in the blocking solution (2 hr at RT or ON at 4°C). Next, embryos were washed four times in PBST (15 min/wash) at RT and mounted before confocal imaging. Antibody pairs and dilutions: chicken anti-GFP (1:1000) and Alexa Fluor 488 goat anti-chicken (1:1000); rat anti-HA (1:500) and Alexa Fluor 647 donkey anti-rat (1:1000); sheep anti-mCherry[191] (1:1000) and Cy3 donkey anti-sheep (1:1000). See the **KEY RESOURCE TABLE**.

### Zebrafish: morpholino injections

We diluted the splice-blocking MO2-*plxnd1* morpholino[59] (GeneTools, LLC) in water with phenol red. We delivered 2.5 ng of the morpholino into the cytoplasm of one-cell stage zebrafish embryos via microinjection of 2-4 nl of the morpholino solution (as in[59]). Consult the **KEY RESOURCE TABLE**.

### Zebrafish: embryo mounting for confocal imaging

We mounted live and fixed, immunofluorescently stained zebrafish embryos on their side by embedding them within a cooling globule of a 0.5% low-melt AquaPor HR GTAC Agarose (national diagnostics) solution in distilled water. Before mounting live embryos, we anesthetized them in zebrafish system water (ZSW) with 0.4 mg/mL of Tricaine (ZSW-0.4T). To collect images for analyses of EC morphometry with the *Vessel Analyzer* app, we mounted the embryos on a Fisherbrand Frosted Microscope Slide, topping the globule with a square cover glass. For other applications, we mounted the fish in an agarose globule on a 35 mm plastic Petri dish (Falcon). Refer to the **KEY RESOURCE TABLE**.

### Zebrafish: confocal microscopy

We took lateral images of the trunk of live and fixed, immunofluorescently stained embryos of the region found dorsal to the yolk extension. We employed a Leica TCS SP8 confocal microscope, the 488, 561, and 647 nm laser lines, and these Leica objectives: 40x /1.10 N.A. HC PL APO W CORR CS2 water immersion (for image analyses using the *Vessel Analyzer* app) and 40x/0.8 N.A. HCX APO L U-V-I water dipping (for everything else). In the images used for quantification, the total depth span of the Z-stack is larger than the axial vessels’ width. The sections below provide further imaging details.

### Zebrafish: imaging and quantification of the DA caliber

We used confocal imaging (pin-hole size: 1; digital zoom: 0.75; z-step size: 1.2 μm; speed: 600 Hz; line averaging: 2x; 16-bit image resolution; format (image size in pixels): 1024×512 (24, 48 hpf) and 1024×704 (72 hpf); 488 or 561 nm laser lines) of live or fixed, immunostained embryos to visualize the vasculature with the *Tg(fli1:EGFP)^y1^*[53] or *(kdrl:mCherry)^y206^*[185] endothelial cytosolically-targeted fluorescent reporters within a region spanning four (at 24 hpf) or three somites (after 24 hpf). We made orthogonal projections from each Z-stack at roughly the midpoint of each somite to create three or four virtual DA cross-sections. We manually traced the DA’s luminal perimeter in each orthogonal projection and then measured the enclosed area. The average of these measurements is the DA caliber reported for each embryo. We processed the images, traced the DA’s luminal perimeter, and automatically extracted the DA’s caliber using FIJI software (versions 1.52n and 1.54f)[196]. We note that we measured the DA caliber using live or fixed embryos. Importantly, our comparisons involve fish under the same conditions as the figure legends note. While we did not determine the impact of fixation on DA caliber at 24 and 32 hpf, we tested its effects at 48 and 72 hpf. We found no significant difference in DA caliber at 48 hpf between live and fixed embryos. However, at 72 hpf, the DA caliber of fixed embryos appears smaller than that of live fish. Look at the **KEY RESOURCE TABLE**.

### Zebrafish: imaging and quantification of the DA’s EC abundance

We utilized confocal imaging (pin-hole size: 1; digital zooms: 1 (24 hpf), 0.75 (32, 48, and 84 hpf); z-step size:1.2 μm; speed: 600 Hz; line averaging: 2x; 16-bit image resolution; format (image size in pixels): 1024×512 (24, 48 hpf), 1024×704 (72 hpf), 1024×768 (84 hpf); 488 nm laser line), of live embryos to visualize the *Tg(fli:nEGFP)^y7^* pan-endothelial nuclear reporter[100] within a DA segment (145.45 μm at 24 hpf, 193.94 μm at 32 hpf, 218.18 μm at 48 hpf, and 266.67 μm at 72 and 84 hpf), employing the maximum intensity projection from each Z-stack to quantify the number of ECs manually. Guided by the regular positioning of the Se vessels (see[58, 72, 218]), we calculated the average somite length for each stage and genotype (except for *plxnd1* mutants, for which we used the WT-like sibs’ values). Average somite lengths: 29-32 μm (24 hpf), 32-36.6 μm (32 hpf), 42-48 μm (48 hpf), 61.4-66.7 μm (72 hpf) and 68-70 μm (84 hpf). Using the appropriate age-specific value, we converted the span of the imaged DA’s segment into somite lengths. To calculate the ratio of DA’s ECs/somite, we divided the DA’s EC counts by the number of somite lengths. We processed the images using FIJI software (versions 1.52n and 1.54f)[196]. Consult the **KEY RESOURCE TABLE**.

### Zebrafish: imaging and quantification of the DA’s endothelial *Tg(klf2a:H2b-EGFP)^ig11^*levels

We employed confocal imaging (pin-hole size: 1; digital zoom: 0.75; z-step size: 1.2 μm; speed: 600 Hz; line averaging: 2x; 16-bit image resolution; format (image size in pixels): 1024 x704; using the 488 nm and 561 nm laser lines sequentially) of live embryos to co-visualize the signals from the *Tg(klf2a:H2b-EGFP)^ig11^* transcriptional *klf2a* nuclear reporter (expressed in the endothelium and other tissues[87]) and the *Tg(kdrl:nls-mCherry)^y173^*blood endothelium nuclear marker[90]. We used the latter to create a binary mask, automatically defining regions of interest specific to the endothelial signals of the former. Subsequently, we aggregated the intensities of the H2b-EGFP signals across the Z-stack and represented them using a color scale. To prepare the graphs, we first calculated the mean fluorescent intensity for each EC. Then, we averaged the mean intensities of the ECs within the DA for each embryo. We processed the images and extracted fluorescent intensities using FIJI software(versions 1.52n and 1.54f)[196]. Check the **KEY RESOURCE TABLE**.

### Zebrafish: imaging and quantification of EC morphometry with the *Vessel Analyzer* app

We used confocal imaging (pin-hole size: 1; digital zoom: 1; z-step size: 0.42 μm; speed: 600 Hz; line averaging: 2x; 16-bit image resolution; format (image size in pixels): 1024×348; voxel size: 0.2841 x 0.2841 x 0.4249 mm^3^; employing the 488 nm and 561 nm laser lines sequentially) to image in live embryos the tight junction ZO-1 fluorescent protein fusion from the *TgBAC(tjp1a-eGFP)^pd1252^* knock-in line[112] and the *Tg(kdrl:mRFP-F)^y286^*blood endothelium’s cell membrane label[117]. The captured dual-channel image files were processed using our in-house developed *Vessel Analyzer* app programmed on MATLAB (R2021a release; MathWorks®)[198].

This tool offers a detailed EC and vessel morphology analysis, allowing for comparisons across stages and genotypes. The app automatically defines the outer and luminal perimeters and the lumen center in planes orthogonal to the vessel’s axis (“cross-sections”) for the user-selected vessel segment (in this case, a DA portion). It then projects the EC junctions in the vessel wall from the lumen center into a cylinder. Next, the software unwraps the cylinder surface into a plane from the 0-degree line. The unwrapped 2D surface facilitates the identification of individual ECs’ perimeters. The software presents two adjacent plane copies to facilitate the segmentation of EC junctions that cross the 0-degree generating line. The user then manually traces a polygon following the junctional signals to provide a skeleton of the cell perimeter. An algorithm then identifies all junction points close to the polygon to refine the cell’s borders. The refined perimeter of each cell is back-projected in its original position into a 3D vessel model to create a 3D model of the cell. A 3D rendering allows visualization of the vessel’s luminal and outer surfaces, along with EC junctions projected into either surface. Alternatively, the model displays the vessel’s luminal surface and the volume of any selected EC. The 3D models of the vessel and its constituent ECs enable their morphometric quantification. <u>At the cellular level, the app’s outputs include</u> <u>the following single-EC morphometry parameters measured at the vessel’s luminal surface</u>. (**1**) **Perimeter**. (**2**) **Length** (axial length connecting the two perimeter points furthest apart along the vessel’s axis). (**3**) **Width** (the transversal length connecting the two most distant perimeter points). (**4**) **Area** (integration of the total surface occupied by the cell). (**5**) **elongation** (the length/width ratio). The software also calculates the (**6**) **cell’s volume** by integrating the cell’s height between the luminal and basal areas. Additionally, the app uses each EC’s mass distribution (assuming uniform density) to calculate the moment of the inertia tensor around the cell’s center of mass. This mass distribution-dependent parameter measures the cell’s resistance to rotation. With this parameter, the app calculates three more measures for each cell. The (**7**) **mean moment of inertia** (the average moment along all directions). The (**8**) **anisotropy of the inertia moment** (which measures how much the cell’s moment of inertia values differ along its principal axes). The (**9**) **alignment angle** or **angle with the vessel’s axis (θ)**, the arc between the cell’s minor principal axis of rotation (associated with cell length), and the vessel’s longitudinal axis. **<u>The software additionally quantifies the vessel’s 3D model</u>.** For each cross-section, the app provides five vascular morphometry outputs. These are as follows. The (**1**) **outer** and (**2**) **luminal perimeters** that circumscribe the endothelial basal and apical surfaces, respectively. The (**3**) **maximum** and (**4**) **minimum luminal diameters.** The (**5**) **caliber** (the luminal area enclosed by the luminal perimeter). <u>Finally, the software provides two measures of variation of vessel</u> <u>morphometry</u>: (**1**) **variability in vessel caliber** (the standard deviation of the latter) and (**2**) **vessel’s tortuosity** (a measure of how much the vessel deviates from being a straight tube), namely the ratio of the distance connecting lumen centers across cross-sections along the vessel segment over the straight-line distance between its endpoints). The *Vessel Analyzer* app automatically saves all morphometric outputs into two Microsoft Excel tables corresponding to EC and vessel data. We used GraphPad Prism (versions 9.3 and 10.1) software for statistical analysis of this data and graph generation. **See the KEY RESOURCE TABLE**.

### Zebrafish: Quantification of heartbeat frequency with the MicroZebraLab system

We anesthetized the embryos in ZSW-0.4T for 5 minutes and fast-washed them in ZSW to rinse away the anesthetic solution. We mounted the embryos on their side by embedding them within a cooling globule of 0.5% low-melt AquaPor HR GTAC Agarose (national diagnostics) solution in distilled water in the center of a 35 mm plastic Petri dish (Falcon). Before imaging, we added ZSW into the Petri dish and placed it in a 28.5°C incubator for 1 hour to enable heartbeat normalization. We used the MicroZebraLab system (Viewpoint Life Sciences) to perform semi-automated measurements of the heartbeat frequency. This software relies on pixel density changes within the region of interest under transmitted light to calculate cardiovascular parameters; see[51]. We collected 1-minute avi format videos of the heart (both ventricle and atrium) with a frame rate of 30 frames per second and a resolution of 640 x 512 pixels using a trinocular compound microscope (SWIFT SW380T, Swift Optical Instruments Inc) equipped with a Siedentopf head, a mechanical stage, an Abbe condenser, and an LED lamp. The videos were collected using the microscope’s 10x DIN (Deutsche Industrie Norm: 160 mm focal tube length) achromatic objective (0.25 NA and 6.54 mm working distance) and a Teledyne FLIR GS3-U3- 41C6 camera (FLIR Integrated Imaging Solutions, Inc) mounted on the microscope’s third eyepiece. The software ran on a Dell Optiplex 3000 SFF computer with a 256 GB SSD drive for data storage. Consult the **KEY RESOURCE TABLE**.

### Mouse: generation of endothelial *Plxnd1* knockout (*Plxnd1^ECKO^*) animals

We crossed *Tie2^Cre/+^* mice[127] with *Plxnd1^flox/flox^* mice[71] and then back-crossed the *Tie2^Cre/+^;Plxnd1^flox/+^*offspring with *Plxnd1^flox/flox^* mice to produce control (*Plxnd1^flox/flox^, Plxnd1^flox/+^* or *Tie2^Cre/+^;Plxnd1^flox/+^*) and endothelial *Plxnd1* knockout (*Tie2^Cre/+^;Plxnd1^flox/flox^*) embryos. See the **KEY RESOURCE TABLE**.

### Mouse: histology and immunohistochemistry

We harvested embryos at embryonic day 11.5 (E11.5) from timed pregnancies, considering the afternoon of the plug date as E0.5. Subsequently, we fixed the embryos in 4% PFA, dehydrated them in ethanol, and processed them for paraffin embedding and transverse sectioning. Following dewaxing, we prepped the sections for immunostaining by performing antigen retrieval, rehydration, and blocking in 5% BSA-PBST (0.1%) for 2-3 hours at room temperature. We then incubated the sections with primary antibodies overnight at 4°C, followed by washing and incubation in diluted secondary antibodies for 1-2 hrs at room temperature. To visualize the nuclei, we used DAPI counterstain (see[74, 128]). The antibody pairs and their respective dilutions were as follows. Monoclonal rabbit anti-ERG1 (1:100) and Alexa Flour goat anti-rabbit 647 (1:350). Monoclonal rat anti-Pecam1 (1:50) and Alexa Flour goat anti-rat 488 (1:350). We performed standard H&E staining for gross histological analysis as described in[128-130]. Check the **KEY RESOURCE TABLE**.

### Mouse: measurement of the DA’s luminal cross-sectional area and perimeter and statistical analyses

We scored 4 to 5 embryos in each group (control and *Plxnd1^ECKO^*). We used images of 3 sections per embryo from the cardiac level to calculate the luminal area and perimeter of both the right and the left DA. We quantified all images with ImageJ 1.53k software[197]. We conducted statistical analyses using the two-tailed Student’s *t*- test. We present the data as mean ± standard error of the mean (SEM). Significant differences, *p* <0.05. We used GraphPad Prism version 9.3.1 (GraphPad Software, USA) for statistical analyses. See the **KEY RESOURCE TABLE**.

### MVNs: fixation and immunofluorescent staining

We fixed the engineered MVNs in 4% paraformaldehyde solution (Thermo Fisher Scientific) for 15 minutes, rinsing them thrice with Phosphate-Buffered Saline (PBS, Thermo Fisher Scientific) for 5 minutes each time. We applied these solutions through the media channels of the microfluidic devices, placing the devices on a rocker during fixation and staining. We simultaneously blocked and permeabilized using a solution of 10% goat serum (Gibco) and 1% Triton X-100 (Fisher Scientific) in PBS for 3 hours at room temperature. For immunofluorescence, we applied a 1:50 dilution of a primary mouse antibody against human CD144/VE-cadherin (BD Pharminogen) in 1% goat serum (Gibco) and 0.2% Triton X-100 in PBS overnight at 4°C. We rinsed the samples for 6 hours using frequent changes of PBS solution. We incubated the samples in a 1:200 dilution of goat anti-mouse IgG1 AlexFluor 647 (Thermo Fisher Scientific) and 300 nM DAPI (Thermo Fisher Scientific) overnight at 4°C. We subsequently rinsed the MVNs by frequent exchanges in PBS for several hours, leaving the MVNs in PBS overnight at 4°C before imaging. Check the **KEY RESOURCE TABLE**.

### MVNs: confocal microscopy for vessel caliber measurements

We added fluorescent dextran (Texas Red, 70,000 MW) to the media channels of microfluidic devices perfused through the MVNs. We acquired confocal images using an LSM 710 confocal microscope (Zeiss) at a resolution of 2.4 pixels/µm and a z-spacing of 5 µm (10x dry objective). We generated maximum intensity projection (MIP) images and performed background subtraction using Fiji software (version 2.9.0/1.54d)[196]. We used AutoTube software[193] to calculate the vascular caliber. We performed MIP image pre-processing via adaptive histogram equalization, illumination correction, and image denoising through Block-Matching and 3D Filtering. We executed tube detection with the Multi-Otsu thresholding. We removed short ramifications (length smaller than 20 pixels) and merged branch points within a 22-pixel radius. Check the **KEY RESOURCE TABLE**.

### MVNs: confocal microscopy for KLF2-GFP expression quantification

We acquired confocal images using an LSM 710 confocal microscope (Zeiss) at a resolution of 1.2 pixels/µm and a z-spacing of 2 µm (20x dry objective). We generated maximum intensity projection (MIP) images using Fiji software (version 2.9.0/1.54d)[196]. We used Matlab (Version R2022b, The MathWorks, Inc.) to calculate a global threshold level for each image, binarize the image by applying the threshold to identify the GFP-positive pixels, and calculate the percentage of the image that is positive for GFP. Consult the **KEY RESOURCE TABLE**.

### MVNs: confocal microscopy for EC area measurements

Following immunostaining, we performed tiling imaging of the MVNs using an LSM 710 confocal microscope (Zeiss) at a resolution of 2.4 pixels/μm and a z-spacing of 1 μm (40x dry objective). We used the maximum intensity projection (MIP) images to calculate cell area using Fiji software (version 2.9.0/1.54d)[196]. We pre-processed the z-stacks by applying Background Subtraction and Median Filtering before creating MIPs. We further processed the MIPs by applying a Gaussian filter. Then, we segmented the cells using the “Morphological Segmentation” plugin[219] of Fiji before computing the area of each segmented cell. We compared these automated measurements against measurements from manual segmentation of cell borders, confirming that the automatic method provides accurate area computations. See the **KEY RESOURCE TABLE**.

## QUANTIFICATION AND STATISTICAL ANALYSES

### Zebrafish

See the legends of **Figures 1-4, S3-S5,** and **S7**. **Mouse:** See **Figure 6** and the “Quantification of the murine DA’s luminal cross-sectional area and perimeter and statistical analyses” section. **MVNs:** See the legend of **Figure 5**.

## SUPPLEMENTAL INFORMATION

### Zebrafish lines generated in this study

Mutant alleles: *plxnd1^GAP1^* (*plxnd1^skt25^*) and *plxnd1^GAP^*^2^ (*plxnd1^skt26^*); see **Table S1**, **Zebrafish genome editing**, and the **KEY RESOURCE TABLE**. Transgenic lines: *Tg(fliep:2xHA-plxnd1)* (full name: *Tg(fliep:2xHA-plxnd1, myl7:EGFP)^skt27^)* and *Tg(fliep:klf2a)* (full name: *Tg(fliep:klf2a-P2A-mCherry-F, myl7:EGFP)^skt28^*). See **Zebrafish constructs for making transgenic lines, Zebrafish transgenesis,** and the **KEY RESOURCE TABLE**.

### Cell lines generated in this study

None.

## Notes

### Competing Interest Statement

The authors have declared no competing interest.

